# Signal peptide-independent secretion of keratin-19 by pancreatic cancer cells

**DOI:** 10.1101/2025.01.18.633717

**Authors:** Philip Moresco, Jonathan P. Kastan, Jung-in Yang, Rishvanth Prabakar, Francesca Minicozzi, Dexter W. Adams, Paolo Cifani, David A. Tuveson, Douglas T. Fearon

## Abstract

The exclusion of T cells causes immune escape of pancreatic ductal adenocarcinoma (PDA). T cell exclusion is mediated by the interaction between CXCR4 on T cells and its ligand, CXCL12, which is complexed to keratin-19 (KRT19) on the surface of PDA cells. KRT19 secretion by PDA cells is essential to this process but is unusual because KRT19 lacks an endoplasmic reticulum (ER)-directing signal peptide (SP). By using biotinylation by an ER-restricted TurboID system and a split-GFP assay in PDA cells, we demonstrate that KRT19 enters the ER via its “head” domain. Additionally, KRT19 is shown to interact with the signal recognition particle and its secretion is sensitive to canonical protein secretion inhibitors. In vivo, mouse tumors formed with ER-TurboID-expressing PDA cells contain biotinylated KRT19. In contrast, keratin-8 (KRT8), which colocalizes with KRT19 on the surface of PDA cells, does not enter the ER. Rather, KRT8 is externalized via secretory autophagy possibly in a complex with KRT19. Thus, despite lacking a classical SP, PDA cells secrete KRT19 to capture CXCL12 and protect against immune attack.

**Significance Statement:** Pancreatic ductal adenocarcinoma (PDA) is resistant to immunotherapy because T cells are excluded from cancer cell nests. Cancer cells capture cancer associated fibroblast sourced CXCL12, which ligates T cell CXCR4, to exclude T cells from cancer cell nests. CXCL12 is captured by cancer cells via the externalization of the normally intracellular intermediate filament keratin-19 (KRT19). We studied the unconventional secretion of KRT19 and found it is secreted by signal peptide independent entry into the endoplasmic reticulum, as well as via secretory autophagy. Thus, PDA externalized immunosuppressive KRT19 through two unconventional means.

## Introduction

Cancer immunotherapy that targets immune checkpoints on T cells has improved patient outcomes for some solid tumors but most carcinomas, including pancreatic ductal adenocarcinoma (PDA), remain refractory to such therapy^1^. This lack of efficacy is most probably a consequence of the exclusion of T cells from cancer cell nests which precludes the effectiveness of checkpoint inhibitors. One mechanism of T cell exclusion occurs through interactions between the chemokine receptor, CXCR4, on T cells with the chemokine, CXCL12, which is principally derived from cancer associated fibroblasts (CAFs). Inhibiting the CXCL12-CXCR4 interaction with the small molecule inhibitor, AMD3100, resulted in intra-tumoral influx of T cells and sensitized tumors to anti-PD-L1 therapy in an autochthonous mouse model of PDA^2^. Additionally, while mouse and human PDA cancer cells do not produce CXCL12, these cells capture CXCL12 on their surface by binding to the externalized intermediate filament protein, keratin-19 (KRT19), to form a “CXCL12-KRT19 coating”^3^. Similar to AMD3100-treated PDA tumors, tumors formed with *Krt19-*edited mouse PDA cells showed spontaneous intratumoral T cell accumulation and responded to anti-PD-1 antibody therapy^3,4^. The T cell infiltration occurring in tumors formed with *Krt19-*edited PDA cells is accompanied by their activation and production of TNFα and IFNγ. These cytokines induce the formation of a CAF population with lower expression of the T cell-excluding CXCL12, but elevation of the T cell-attracting chemokine, CXCL9, to further enhance the tumor immune reaction^5^. Furthermore, *Krt19-*edited mouse PDA tumors reprogram the tumor microenvironment by recruiting NKT cells which facilitate type I IFN production^6^, an innate cytokine required for tumor control^7^. The relevance of these findings in mouse models of PDA to human disease was shown by studies in patients with PDA and colorectal carcinoma in which inhibition of CXCR4 by AMD3100 showed increased intra-tumoral infiltration and activation of T cells^8–10^. In summary, reports from our group^2,3,5,6,8^ and others^4,9,10^ highlights the CXCL12-KRT19 complex as a means by which tumors can evade immune control.

KRT19, a member of the type I intermediate filament family of proteins, heterodimerizes and polymerizes with its type II partner, KRT8, to form part of the cytoskeleton in epithelial cells and carcinomas^11^. KRT19 differs from other type I keratins by its lack of a tail domain and by having a head domain with a unique sequence. Despite these unusual structural features, a germline knockout of *Krt19* alone results in no abnormal phenotype^12^, because perhaps KRT18, which is co-expressed in epithelial cells, also heteropolymerizes with KRT8. Indeed, a phenotype for *Krt19^-/-^* mice is only unmasked when these mice are bred in a *Krt18^-/-^* background^13^. An alternative explanation for *Krt19^-/-^*mice lacking an overt phenotype is the possibility that its role is primarily through immune regulation, as deletion of mouse immune genes often does not result in abnormal development under non-perturbed conditions but only in association with immunological challenge. In accordance with this principle, a role for the specific, saturable binding of CXCL12 by KRT19 was seen only when tumors were formed with sgKRT19-edited PDA cells, resulting in enhanced T cell infiltration and diminished growth^3^.

For KRT19 to interact with T cells as a component of the “CXCL12-coat” on cancer cells, this intermediate filament protein must be secreted. It is noteworthy, despite groups suggesting that the KRT19 secretion occurs^3,14,15^, KRT19 does not have a canonical endoplasmic reticulum (ER)-targeting signal peptide (SP). Proteins that are secreted tend to follow one of three general mechanisms for their externalization. First, the majority of secreted proteins are externalized through the canonical ER-Golgi pathway. These secreted proteins begin their translation in the cytosol, and once their N-terminal SP is translated, the nascent polypeptide interacts with the signal recognition particle (SRP)^16^. The SRP:SP:ribosome complex then binds with the SRP receptor on the ER surface^16^. The SRP receptor interacts with the Sec61 translocon, and the Sec61 complex facilitates protein entry into the ER lumen^17–19^. The second most studied mechanism of protein secretion is unconventional protein secretion (UcPS). UcPS represents a diverse group of pathways that mediate the externalization of cytosolic proteins which lack an ER-targeting SP, or proteins that enter the ER through a SP, but then bypass the Golgi^20^. Some of the pathways of UcPS of SP-lacking cytosolic proteins include the pore mediated pathways (i.e. GSDMD and IL-1β)^21^, direct translocation across the cell membrane (i.e. FGF2)^22^, and vesicle-based secretion pathways (i.e. secretary autophagy and exosome biogenesis)^23,24^. Third, a rare subset of proteins that do not contain an ER-targeting SP have been reported to enter^25^ and be externalized via^26^ the ER or ER-Golgi intermediate complex^27^.

Determining how KRT19 is externalized will provide mechanistic insight into how cancer cells protect themselves from immune attack and is essential for understanding tumor immune suppression. Therefore, we set out to describe the pathways through which KRT19 secretion occurs.

## Results

### Detection of KRT19 on the Plasma Membrane and within the Endoplasmic Reticulum of PDA Cells

Intact FC1242 cells, derived from LSL-KrasG12D/+; LSL-Trp53R172H/+; Pdx1-Cre (KPC) mouse PDA tumors^28^, were stained prior to fixation and permeabilization with monoclonal antibodies specific for KRT19 and KRT8 and examined by confocal microscopy. Images showed these proteins forming filament-like networks that were reminiscent of intracellular intermediate filaments (**Figure 1A**), similar to our previous report^3^. Additional evidence for this extracellular location of KRT19 and KRT8 was provided when cells were labeled with the cell-impermeable sulfo-NHS-LC-biotin and proteins were isolated by streptavidin-pulldown. Immunoblot analysis showed that KRT19 and KRT8 bound to the streptavidin under stringent wash conditions, implying their surface biotinylation similar to a known plasma membrane protein, E-cadherin. An intracellular protein, GAPDH, that served to control for the integrity of the cells, did not bind to the streptavidin and, thus, was not biotinylated (**Figure S1**).

**Figure 1:**
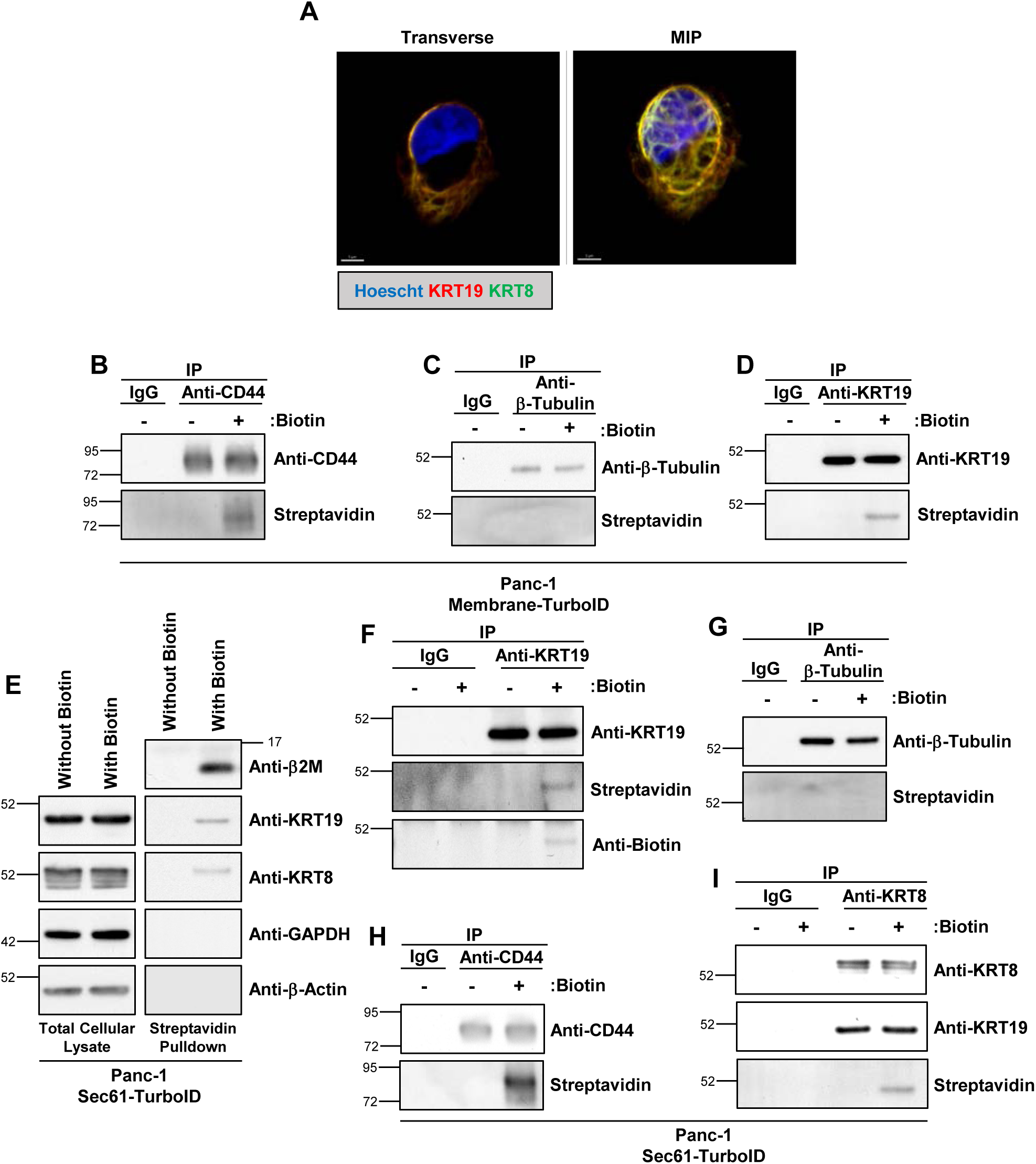
ER resident Sec61-TurboID and biotinylation of KRT19. **(A)** FC1242 cells were stained prior to fixation and permeabilization with antibodies to KRT19 and KRT8 and z-stacks were taken of stained cells. (Scale = 5 μm. Ex **(B-D)** Panc-1 cells expressing membrane-TurboID were cultured in the presence or absence of 50 μM biotin for 18hrs and then lysed with RIPA buffer. The cell lysates were incubated overnight with antibodies to CD44 **(B)**, β-Tubulin **(C)**, and KRT19 **(D)**, respectively, or IgG control. The immunoprecipitated proteins were analyzed by immunoblot for biotin incorporation. **(E)** Panc-1 Sec61-TurboID cells were cultured in the presence or absence of 50 μM biotin for 18hrs and then lysed with RIPA buffer. The cell lysates were incubated with streptavidin-agarose beads to isolate biotinylated proteins. Total cellular lysates and streptavidin-pulldowns were analyzed by immunoblots. **(F-I)** Panc-1 Sec61-TurboID cells were cultured in the presence or absence of 50 μM biotin for 18hrs and lysed with RIPA buffer. One mg/mL lysate was incubated with antibodies to KRT19 **(F)**, β-Tubulin **(G)**, CD44 **(H)**, and KRT8 **(I)**, respectively, or IgG control overnight. The cell lysates and immunoprecipitated proteins were analyzed by immunoblots. Shown are representative results of three replicate experiments. MIP=mean intensity projection

The localization of KRT19 in the secretory pathway in cancer cells was probed by expressing a promiscuous biotin ligase, TurboID, in human Panc-1 cells as a fusion protein with the transmembrane domain of PDGFRβ (TM-TurboID). This fusion protein localizes TurboID to the ER-Golgi lumen and the extracellular space (**Figure 1B-D**)^29^. As positive and negative controls, respectively, an anti-CD44 antibody immunoprecipitated biotinylated CD44 (**Figure 1B**), but an anti-β-tubulin antibody immunoprecipitation (IP) did not reveal biotinylated β-tubulin (**Figure 1C**). Performing an anti-KRT19 antibody IP from lysates of biotin-treated Panc-1 cells expressing the TM-TurboID allowed the detection of biotinylated KRT19 by blotting with streptavidin (**Figure 1D**), providing another line of evidence for keratin secretion.

These data extend our previous report^3^ that showed KRT19 and KRT8 were externalized by PDA cells, and prompted an analysis of the cellular pathway of secretion. To study the potential mechanism of KRT19 secretion, we performed mass spectrometry of the proteins co-immunoprecipitating with KRT19 from the detergent lysates of FC1242 PDA cells. The proteins were resolved by SDS-PAGE and a silver stain depicted an approximately 70 kDa protein band, which was absent from the IgG control IP (**Figure S2A**). LC-MS of this band resolved three protein-folding chaperones, two of which were the HSPA8 and HSAP9 proteins that are present in the cytoplasm and mitochondria, respectively, and the third was HSPA5 (BiP), an ER-resident chaperone that facilitates protein folding and unidirectional protein entry into the ER (**Figure S2B**; **Table S1**)^30^. We confirmed that KRT19 interacts with BiP by observing that BIP co-immunoprecipitates with KRT19 from lysates of both FC1242 cells and human Panc-1 PDA cells (**Figure S2C**). These findings suggested that KRT19 may enter the ER, despite the absence of a SP.

Canonically secreted proteins enter the ER through a multiprotein translocon comprised of Sec61α/β/γ^18,19^. We asked whether KRT19 enters the ER by this mechanism by using Sec61β−tagged with TurboID (Sec61-TurboID), which localizes TurboID within the ER-lumen^31^. To verify the appropriate cellular distribution of Sec61-TurboID, we fractionated Panc-1 cells expressing this fusion protein with digitonin to isolate the cytosolic and cell membrane fractions, and the latter was subjected to n-Dodecyl-β-D-maltoside (DDM) to isolate the ER fraction^32^. Sec61-TurboID was found only in the DDM-solubilized ER fraction, along with endogenous Sec61β and the ER chaperone BiP, whereas GAPDH was enriched in the cytosolic fraction (**Figure S3A**). Furthermore, we found that culturing cells with biotin led to immunofluorescent staining of the incorporated biotin co-localizing with GRP94, an ER marker, and the Golgi marker protein, GM130 (**Figure S3B**), indicating biotinylated proteins were appropriately localizing to the secretory compartment of Sec61-TurboID-expressing cells. We next analyzed the Sec61-TurboID-dependent secretome common to both Panc-1 and BXPC3 human PDA cells. Here we treated Panc-1 and BXPC3 cells expressing Sec61-TurboID and their parental counterparts with biotin, collected the supernatants, isolated biotinylated proteins by streptavidin purification, and then performed LC-MS of the isolated proteins. Of the 63 proteins commonly enriched (Log_2_ fold change > 3.5) in the Sec61-TuboID-expressing Panc-1 and BXPC3 pulldowns, 62 (98.4%) had either a predicted signal peptide or transmembrane domain (**Figure S3C-D**). Together, these data highlight the appropriate distribution and specificity of the Sec61-TurboID system in labeling ER-derived proteins.

With the evidence that the Sec61-TurboID construct was appropriately targeted to the ER, we next sought to determine if KRT19 was biotinylated by Sec61-TurboID. We treated Sec61-TurboID Panc-1 cells with biotin, performed a streptavidin-pulldown from cell lysates, and resolved the proteins by a reducing SDS-PAGE. Immunoblots identified the presence of β2M, ITGB1, Met, PTK7, and GRN, which served as positive controls. GAPDH, β-actin, ANXA1, and ERK1/2 served as the negative controls and were absent from the pulldown proteins. Accordingly, the identification in the streptavidin-pulldown from biotin-treated Sec61-TurboID PDA cells of KRT19 and its binding partner, KRT8 (**Figure 1E; Figure S3E**) suggests either or both proteins may enter the ER without a SP.

The streptavidin-pulldown of biotin-treated Sec61-TurboID cell lysates can result in the isolation of biotinylated proteins or non-biotinylated proteins that interact with biotinylated binding partners. To determine if KRT19 is directly biotinylated in the Sec61-TurboID system, we cultured cells in the presence or absence of biotin, performed an anti-KRT19 antibody IP from the cell lysates, and blotted with anti-biotin antibody and streptavidin-HRP, respectively. Both anti-biotin antibody and streptavidin-HRP identified a biotinylated protein having the molecular weight of KRT19 in the anti-KRT19 antibody IP (**Figure 1F**). The same pattern was seen with IPs targeting CD44, a *bona fide* secretory protein, while IPs targeting β−tubulin did not isolate a biotinylated protein (**Figure 1G-H**). Repeating these experiments with an anti-KRT8 antibody IP resulted in the identification of a biotinylated protein the molecular weight of KRT19, but not that of KRT8. Consistent with this finding, anti-KRT19 antibody immunoblot of the KRT8 IP revealed KRT19, demonstrating that KRT8 and biotinylated KRT19 had formed a complex (**Figure 1I**). We confirmed that biotinylation of KRT19 depended on the presence of Sec61-TurboID and was not a consequence of an endogenous biotinylation reaction by culturing both the parental and the Sec61-TurboID-expressing Panc-1 cells with exogenous biotin and then immunoprecipitating KRT19; an anti-CD44 IP served as a positive control. Similar to the anti-CD44 IP, antibody to KRT19 isolated biotinylated KRT19 from Sec61-TurboID expressing, but not the parental, Panc-1 cells (**Figure S4A**). We sought to expand these results to include mouse PDA cells and found KRT19 is similarly biotinylated in FC1242 cells ectopically expressing Sec61-TurboID, along with β2M, but not β-tubulin (**Figure S4B-D**). Therefore, KRT19 is biotinylated by the Sec61-TurboID construct, in contrast to KRT8 and the cytoplasmic proteins GAPDH, β-actin, and β-tubulin.

As an alternative method to evaluate KRT19 entry into the ER, we adapted a split-GFP reporter assay^33^. Here, the S1-10 fragment of GFP is expressed in the ER by fusion with a SP and ER-retention tag, or S1-10 is expressed in the cytosol by removal of the SP and ER-retention tag. The complementing GFP-S11-fragment, which combines with the S1-10 fragment to emit GFP fluorescence, is fused to the N- or C-terminus of KRT19, KRT18, and KRT8 to assess whether these proteins can enter the ER (**Figure 2A**). SP-bearing S11-KRT19 (SP-S11-KRT19) served as the positive control for protein entry into the ER. Transient transfection of HEK293T cells with ER-localized S1-10 showed that when cells were co-transfected with KRT19-S11, but interestingly not S11-KRT19, nor any combination of KRT18 and KRT8 GFP-S11 fusion proteins, a GFP-signal was seen by flow cytometry (**Figure 2B**; **Figure S5**). In contrast, all combinations of S11 N- or C-terminal fusions with KRT19, KRT18, and KRT8 resulted in GFP signal only when the S1-10 GFP fragment was expressed in the cytosol (**Figure S5**). Thus, in an entirely orthogonal assay, KRT19 is shown to enter the ER.

**Figure 2:**
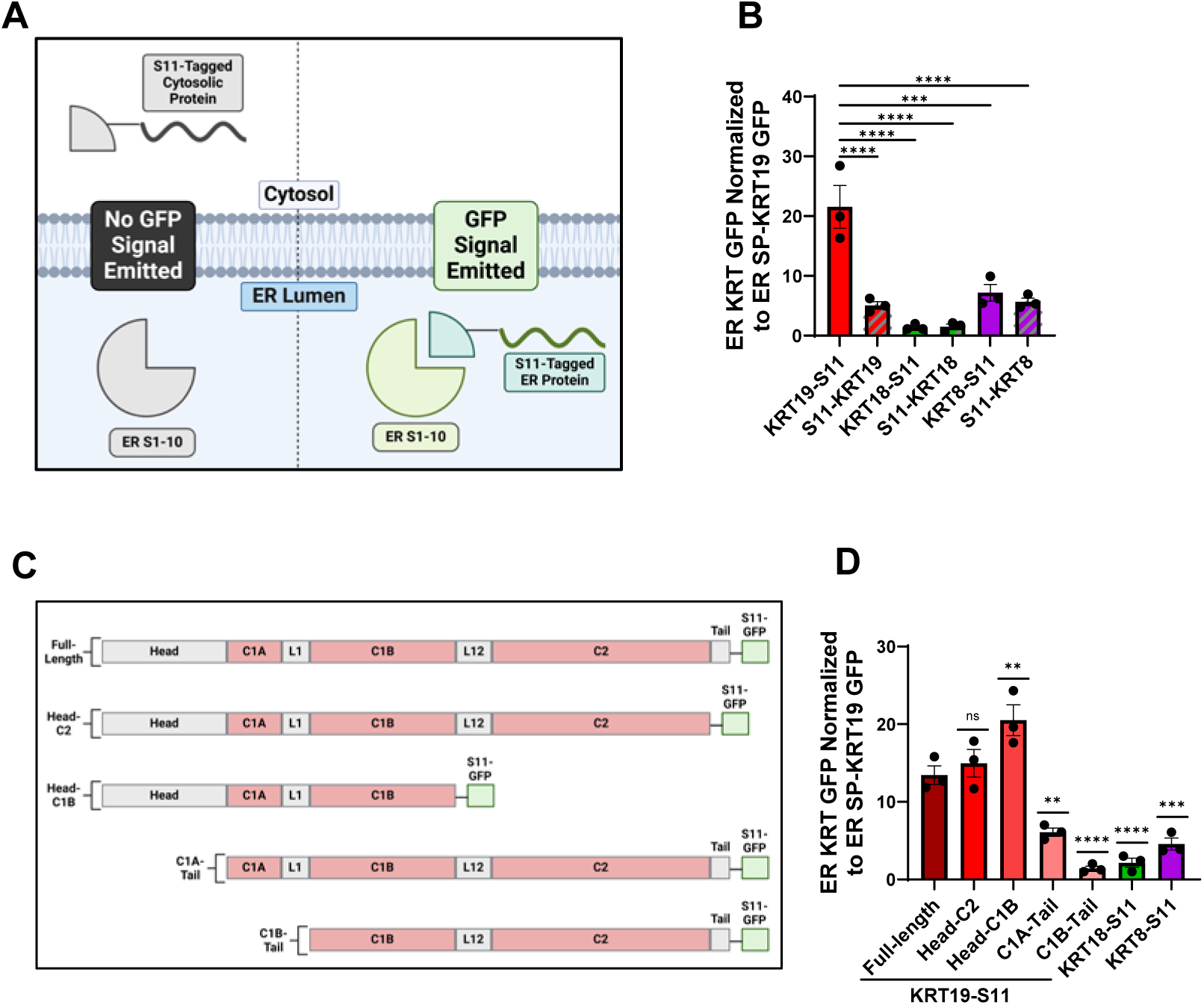
The Split-GFP assay of KRT19 entry into the ER. **(A)** Schematic of the Split-GFP reporter assay. **(B)** The GFP signal obtained from cells expressing ER-targeted GFP domains S1-10 and KRT8, KRT18 or KRT19 fused to the N-terminus or C-terminus of GFP-domain S11 was normalized by dividing the percent GFP-positive cells that expressed the ER-targeted GFP-domains S1-10 and signal peptide (SP)-modified GFP-domain S11-KRT19 fusion protein. **(C)** Schematic of the KRT19 truncations for Split-GFP reporter assay. **(D)** The GFP signal obtained from cells expressing ER-targeted GFP-domain S1-10 and the KRT19 truncations fused to GFP-domain S11 was normalized by dividing the percent GFP positive cells expressing ER S1-10 and SP-S11-KRT19. Shown are representative results of three replicate experiments. Mean ±SEM is plotted; ns=not significant; **p<0.01, ***p<0.001; ****p<0.0001; SP=signal peptide.

The absence of ER entry of N-terminally tagged S11-KRT19 suggested that the N-terminus of KRT19 may provide an important signal to its ER-trafficking. To identify more definitively the region of KRT19 that is required for entry into the ER, we expressed truncated forms of KRT19 fused at their C-termini with the GFP-S11 fragment in cells expressing either cytosolic or ER-resident S1-10 (**Figure 2C**). Removing the N-terminal head domain of KRT19 abrogated the ability of the KRT19-S11 fusion protein to fluoresce in cells expressing ER-localized S1-10 (**Figure 2D; Figure S6**) and is consistent with the head domain of KRT19 functionally serving as its SP. All KRT19 truncations fused with S11 produced GFP signal only when the S1-10 fragment of GFP was expressed in the cytosol (**Figure S6**). An alternative possibility that KRT19 entry into the ER was mediated by a gene translocation that appends a SP onto the coding region of KRT19 was excluded by long-read RNA sequencing, as all the reads of *KRT19* transcripts mapped to its exonic sequences (**Figure S7**). Therefore, KRT19 traffics to the ER and its head domain is required for this process.

### KRT19 Secretion is Sensitive to Inhibitors of ER-Golgi Trafficking

To examine KRT19 secretion by individual PDA cells, we adapted a KRT19 enzyme-linked immunosorbent spot (ELISpot) assay^14^. In this assay, PDA cells are grown in wells precoated with a mouse anti-KRT19 capture antibody targeting coil 2 of KRT19 and, after removing the cells, the externalized KRT19 bound to the capture antibody is revealed by incubation with a rabbit anti-KRT19 detection antibody. We performed the ELISpot assays with two anti-KRT19 detection antibodies targeting the head and the tail of KRT19, respectively, to validate that full-length KRT19 was secreted rather than the caspase 3-cleaved KRT19 fragment that is released by apoptotic cells (**Figure 3A**; **Figure S8A-B**)^34^. When performed with human Panc-1 cells, this assay detects secretion of full-length KRT19. As a control, we observed that ELISpot puncta were lost when the anti-KRT19 capture antibody was replaced with a non-immune IgG control. Furthermore, when the capture and detection anti-KRT19 antibodies were replaced with anti-ERK1 antibodies, ERK1 ELISpot puncta did not appear (**Figure S8C-D**).

**Figure 3:**
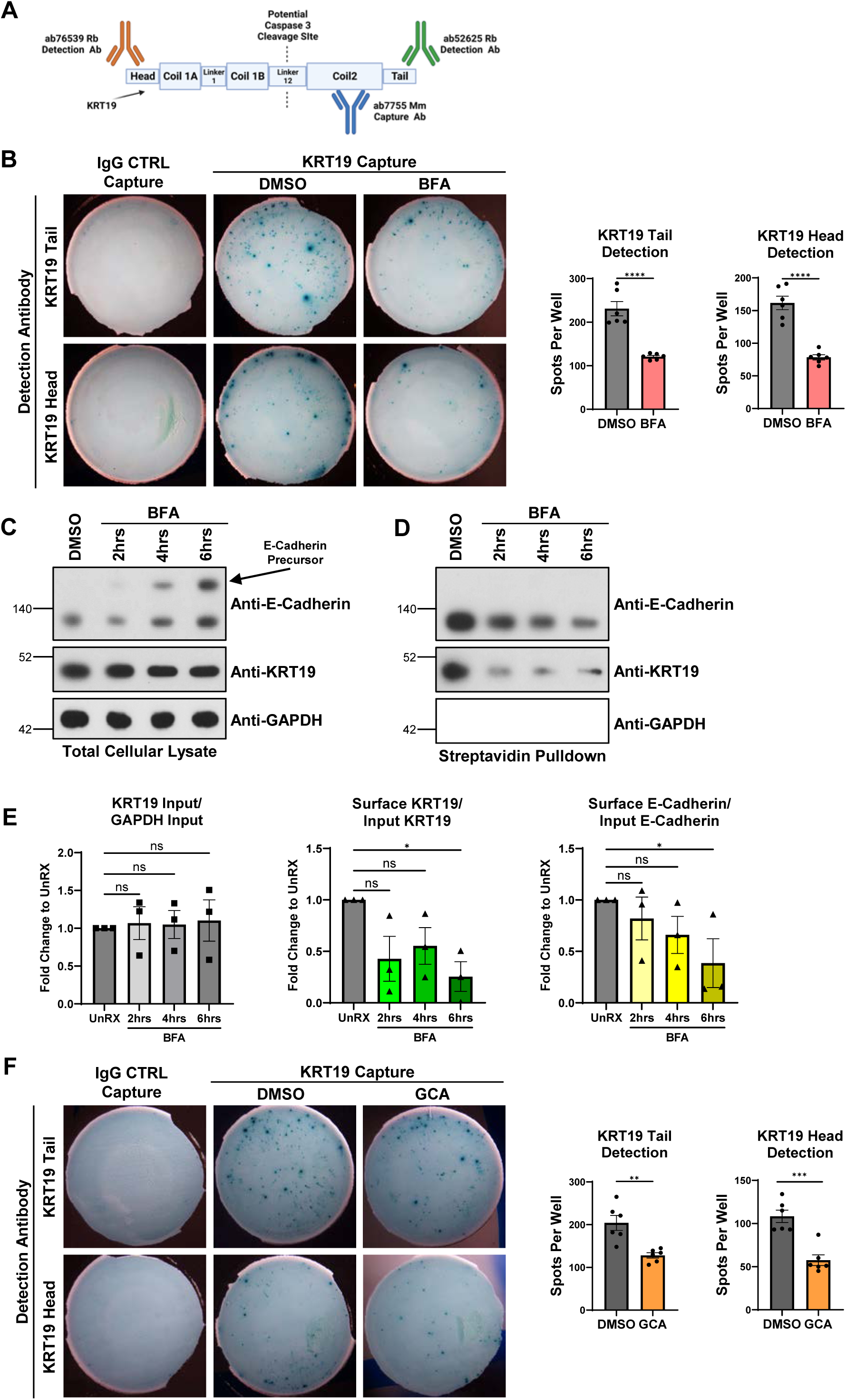
Modulation of KRT19 secretion via BFA and GCA. **(A)** Schematic of the anti-KRT19 antibodies used for the ELISpot assays. **(B)** Panc-1 cells, 1000, were cultured in the presence of DMSO or 11 μM BFA for 18 hrs in ELISpot wells that were coated with CTRL IgG and the anti-KRT19 capture antibody. The captured KRT19 was detected by anti-KRT19 antibodies that were specific for the head and tail domain, respectively. Shown are representative results of two replicate experiments with three replicate wells per experiment. A Student’s T test was performed. **(C-E)** Immunoblots are shown of total cellular lysates (**C**) or cell surface proteins **(D)** isolated from PDA FC1242 cells. **(E)** The fold-change in total KRT19, KRT19 secretion, and E-cadherin secretion occurring with 1.25 μM BFA treatment for two, four, or six hours. One-way ANOVA with Dunn’s multiple comparisons was performed. A representative experiment of three independent experiments is shown. **(C)** Panc-1 cells, 1000, were cultured in the presence of DMSO or 10 μM GCA for 12 hrs in ELISpot wells that were coated with CTRL IgG and anti-KRT19 capture antibody. The captured KRT19 was detected by anti-KRT19 antibodies that were specific for the head and tail domain, respectively. Shown are representative results of two replicate experiments with three replicate wells per experiment. A Student’s T test was performed. Mean±SEM is plotted; CTRL=control; ns=not significant; *p<0.05, **p<0.01, ***p<0.001; ****p<0.0001

We assessed the effects of treating Panc-1 cells with Brefeldin A (BFA), a fungal macrolide that inhibits the trafficking of secretory vesicles from the ER to the Golgi. Consistent with the biotin-labeling of KRT19 in the Panc-1 cells expressing Sec61-TurboID, significantly fewer KRT19 ELISpot puncta were generated by the Panc-1 cells when they were treated with BFA (**Figure 3B**). Total KRT19 levels were unchanged in BFA-treated cells, and BFA did not affect cell viability (**Figure S8E-F**). Furthermore, treating FC1242 cells with BFA for two, four, and six hours resulted in decreased KRT19 secretion. This decrease in KRT19 secretion mirrored that of E-cadherin, and did not affect total KRT19 levels within the cell (**Figure 3C-E)**. Consistent with this indication that KRT19 was secreted by trafficking through the Golgi, treatment of Panc-1 cells with Golgicide A (GCA), a specific inhibitor of GBF1 that blocks ER to Golgi protein transport, also significantly decreased KRT19 ELISpot puncta (**Figure 3C**). GCA did not affect total KRT19 levels or Panc-1 cell viability (**Figure S8E-H)**. Thus, a portion of secreted KRT19 by Panc-1 cells is through the ER-Golgi pathway.

### KRT19 Interacts with the SRP and Sec61

To assess whether KRT19 enters the ER as a result of interacting with proteins that mediate secretion of SP-containing proteins, we assessed whether KRT19 interacts with the SRP, the cytosolic ribonucleoprotein which brings the translating ribosome to the ER surface^16,35^. Immunoprecipitated proteins obtained with anti-SRP68 antibody and an anti-SRP19 antibody, respectively, from lysates of Panc-1 cells demonstrated that both SRP68 and SRP19 co-precipitated KRT19, but not GAPDH (**Figure 4A-B**). We sought to determine whether KRT19 also interacted with Sec61, or V5-tagged Sec61, by performing IPs with an anti-Sec61β antibody and an anti-V5 antibody. KRT19 coprecipitated with both antibodies (**Figure 4C**). An anti-Sec61β antibody also co-precipitated KRT19 from parental Panc-1 cells, excluding an effect of the TurboID (**Figure 4D**). Taken together, these data show that KRT19 interacts with the biochemical machinery required for traditional protein entry into the ER, despite lacking a canonical SP.

**Figure 4:**
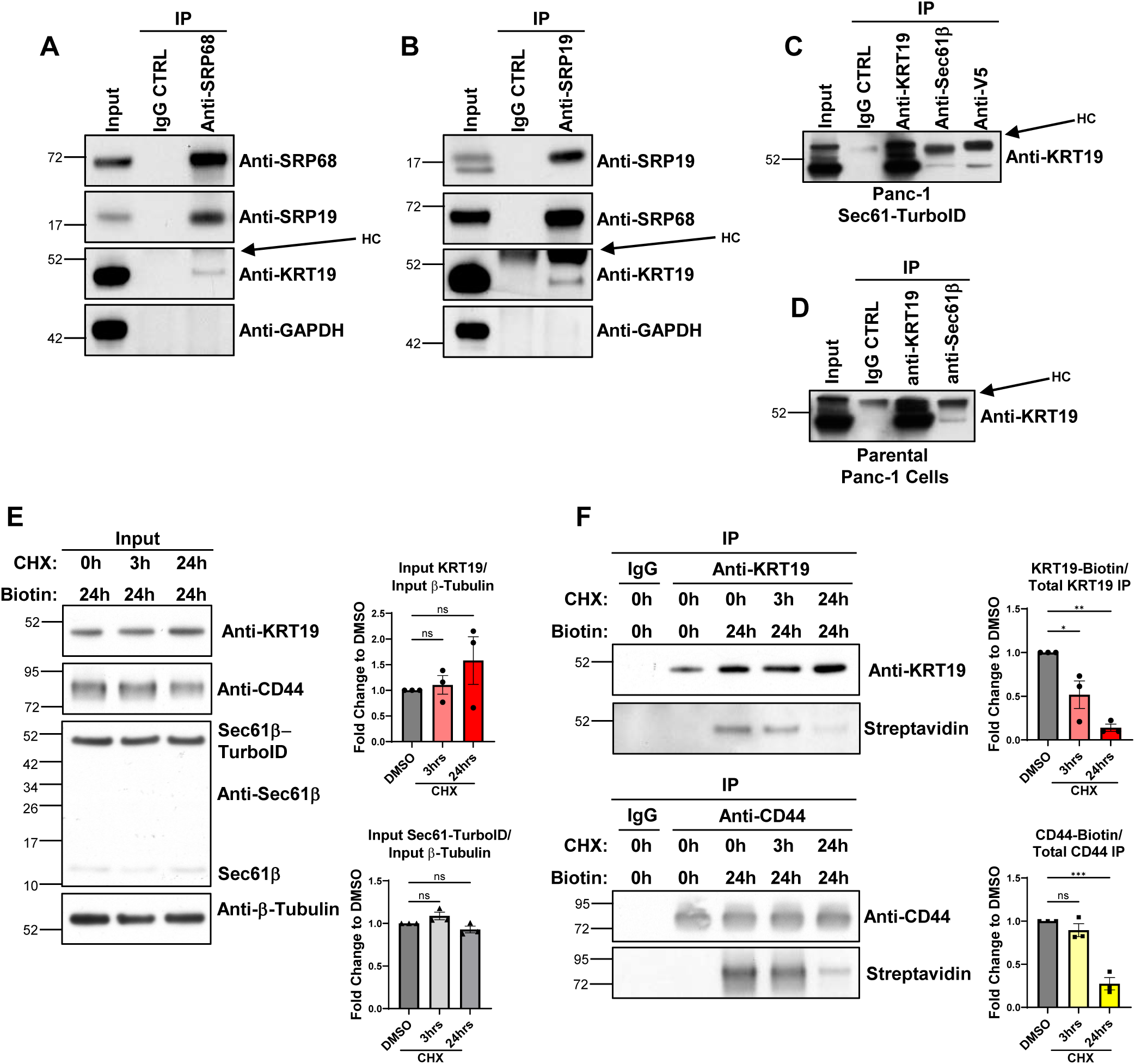
KRT19 cotranslational entry into the ER. **(A, B)** Panc-1 cells expressing Sec61-TurboID were lysed with 1% TritonX-100. The lysate was incubated overnight with antibodies to SRP68 **(A)**, SRP19 **(B)**, and IgG CTRL, respectively. Cell lysates and the immunoprecipitated proteins were assessed by immunoblots. Shown are representative results of three replicate experiments. **(C, D)** Panc1 cells expressing Sec61-TurboID **(C)** or parental Panc-1 cells **(D)** were lysed with 1% TritonX-100 and lysates were incubated overnight with antibodies to KRT19, Sec61β, V5, or IgG CTRL. Cell lysates and the immunoprecipitated proteins were assessed by immunoblots. Shown are representative results of three replicate experiments. **(E-F)** Panc-1 cells expressing Sec61-TurboID cells were cultured for 24 hrs in the presence or absence of 50 μM biotin, and for timed intervals with 50 μg/mL CHX. Lysates of the cells that had been obtained with RIPA buffer were incubated overnight with antibodies to KRT19, CD44, and IgG CTRL. Total lysates **(E)** and immunoprecipitated proteins **(F)** were assessed by immunoblots. Shown are representative results of three replicate experiments. A one-way ANOVA (Brown-Forsythe test) with Dunnett’s multiple comparisons was performed. Mean ± SEM is plotted; h=hours; ns=not significant; *p<0.05, **p<0.01; ***p<0.001; ****p<0.0001; HC=heavy chain.

### Krt19 Enters the ER for Co-translational Secretion

Proteins typically enter the ER co-translationally^36^. We performed a cycloheximide (CHX) pulse experiment with continuous biotin treatment to determine if the biotinylation of KRT19 by Sec61-TurboID is a co-translational process. We found that three hours or 24 hours of CHX treatment did not affect total KRT19 levels (**Figure 4E**), but did suppress incorporation of biotin into KRT19 (**Figure 4F**). We demonstrated that the recovery of biotinylated CD44 was similarly affected by CHX treatment (**Figure 4F**) and that Sec61-TurboID levels were unchanged by CHX treatment (**Figure 4E**), excluding an alternative explanation for the decreased biotinylated KRT19 and CD44. We also sought to determine if KRT19 secretion was dependent upon protein translation. With the ELISpot assay of KRT19 secretion, we observed that, while the number of KRT19 puncta was not decreased with CHX addition, the size of these puncta were smaller when either KRT19 detection antibody was used (**Figure S9**). Thus, newly synthesized KRT19, not pre-existing cytoplasmic KRT19, traverses the Sec61 complex to enter the ER for its ultimate secretion.

### KRT19 Enters the ER and is Secreted Through the Sec61 Translocon

To confirm that KRT19 traverses the Sec61 translocon, as suggested by biotinylation via Sec61-TurboID, we treated Sec61-TurboID expressing Panc-1 cells with Eeyarestatin I (ESI), a Sec61 pore blocking agent^37^. ESI treatment of the Panc-1 cells expressing Sec61-TurboID diminished the biotinylation of KRT19 and CD44 (**Figure 5A-B**), both after six hours (**Figure S10B**) and 18 hours (**Figure 5B**) of treatment. The decreased biotinylation of KRT19 could not be ascribed to lower levels of Sec61-TurboID or total KRT19 (**Figure 5A**; **Figure S10A**). As ESI also inhibits p97, which is involved in ER associated protein degradation (ERAD)^38^, we determined if the ESI-mediated decrease in KRT19 biotinylation results from ERAD inhibition. However, treatment of Sec61-TurboID expressing Panc-1 cells with CB-5083, a selective p97 inhibitor^39^, did not affect KRT19 biotinylation (**Figure S10C-D**). Finally, treating Panc-1 cells with ESI in the ELISpot assay decreased KRT19 secretion (**Figure 5C**), demonstrating that a portion of KRT19 enters the ER via Sec61 for its secretion.

**Figure 5:**
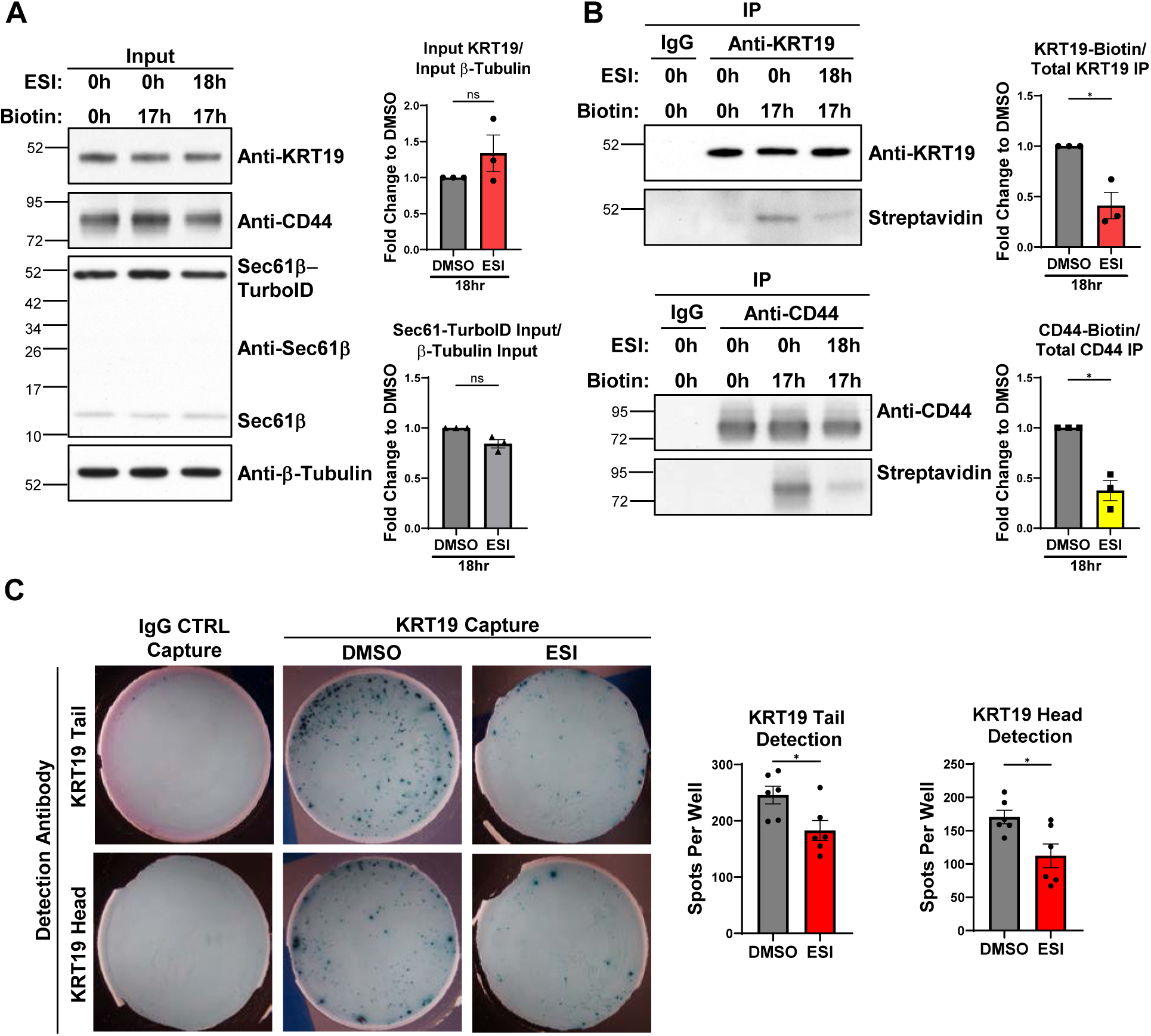
Entry of KRT19 into the ER *via* Sec61. **(A-B)** Panc-1 cells expressing Sec61-TurboID were pre-treated for 1 hr with 10 μM ESI, and then cultured for 17hrs in the presence or absence of ESI and 50 μM biotin. Lysates of the cells that had been obtained with RIPA buffer were incubated overnight with antibodies to KRT19, CD44, and IgG CTRL, respectively, and total lysates **(A)** and the immunoprecipitated proteins **(B)** were assessed by immunoblots. Shown are representative results of three replicate experiments. Welch’s T test was performed. **(C)** Panc-1 cells, 1000, were cultured in the presence of DMSO or 10 μM ESI for 18 hrs in ELISpot wells that were coated with CTRL IgG and anti-KRT19 capture antibody. The captured KRT19 was detected by anti-KRT19 antibodies that were specific for the head and tail domain, respectively. Shown are representative results of three replicate experiments with two replicate wells per experiment. A Student’s T test was performed. Mean ± SEM is plotted; h=hours; ns=not significant; CTRL=control; *p<0.05.

### Secretory Autophagy of KRT19 and KRT8

Inhibition, but not elimination, of KRT19 secretion by BFA, GCA, CHX, and ESI suggested that another means of KRT19 secretion may exist. Secretory autophagy is one pathway of UcPS and also has been shown to be immune suppressive in PDA^20,40^. To evaluate if secretory autophagy has a role in KRT19 secretion, we expressed a doxycycline-inducible dominant negative inhibitor of autophagy, mSt-ATG4B^C74A^, in mouse FC1242 PDA cells. mSt-ATG4B^C74A^ inhibits autophagy by sequestering LC3 paralogues, blocking the formation of the Atg7-LC3 intermediate and preventing autophagosome closure^41^. Inducing mSt-ATG4B^C74A^ decreased KRT19 secretion in the ELISpot assay (**Figure 6A**) without affecting total KRT19 levels (**Figure S11A**). Autophagy and the canonical Golgi-mediated mechanism were independent secretory pathways for KRT19, as BFA treatment, at a concentration did not affect cell viability or total KRT19 levels (**Figure S11B-C**), decreased KRT19 secretion in mouse PDA FC1242 cells (**Figure 6B**), and also decreased KRT19 secretion from doxycycline-treated FC1242 cells expressing mSt-ATG4B^C74A^ (**Figure 6B**).

**Figure 6:**
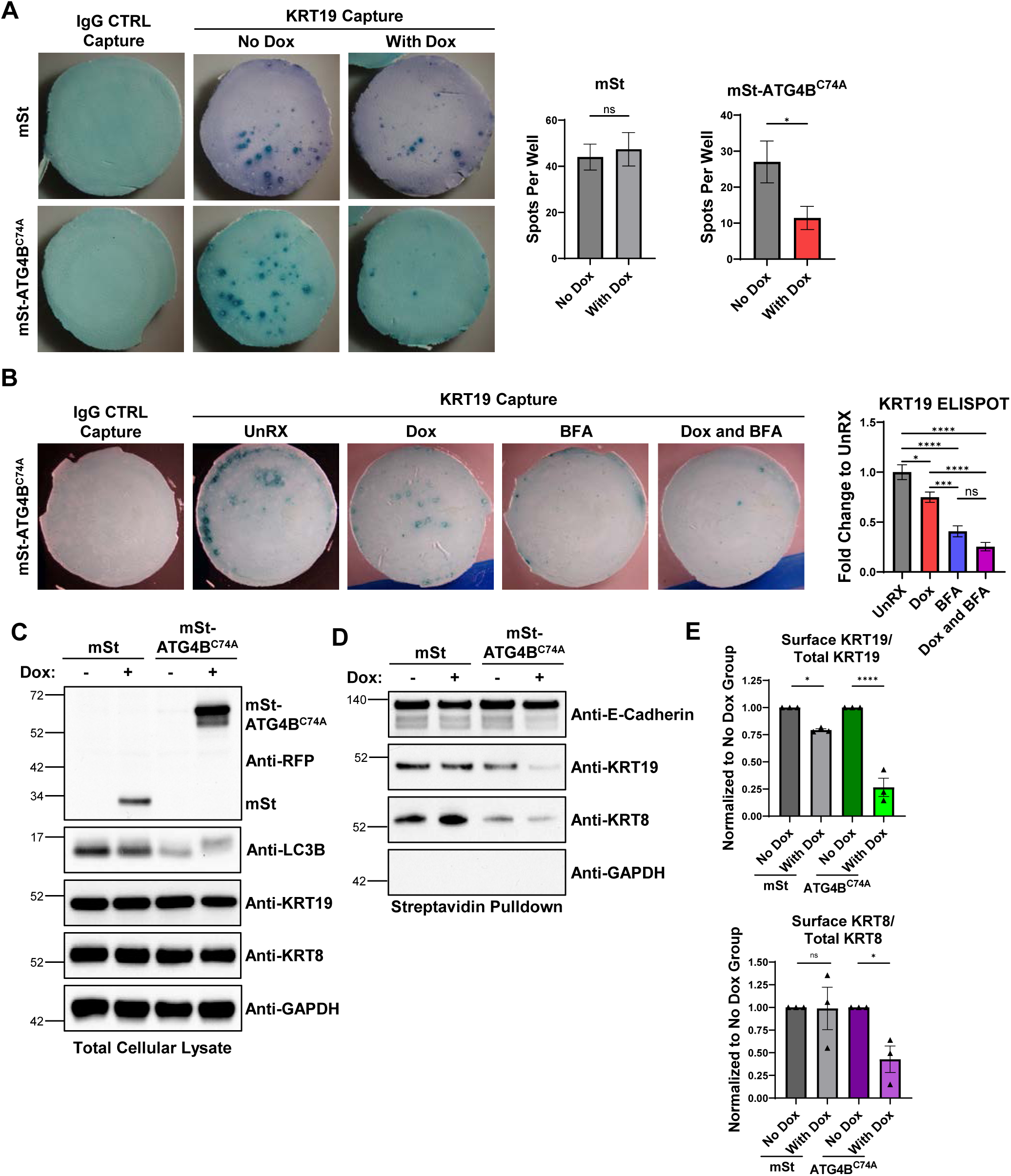
Secretion of KRT8 and KRT19 through autophagy. **(A)** Mouse PDA FC1242 cells, 1000, expressing doxycycline-inducible mSt or mSt-ATG4B^C74A^ were plated in ELISpot wells coated with IgG CTRL or anti-KRT19 capture antibody. The cells were cultured for 48 hrs in the presence or absence of 1 μg/mL doxycycline. Antibody specific for KRT19 tail was used for detection. A representative experiment of three independent experiments is shown with two to three replicate wells per experiment. A student’s T test was performed. **(B)** 1500 mouse PDA FC1242 cells expressing doxycycline-inducible mSt-ATG4B^C74A^ were plated in ELISPOT wells coated with non-specific IgG (CTRL) or anti-KRT19 capture antibody. The cells were cultured for 48 hrs in the presence or absence of 1 μg/mL doxycycline; DMSO or 1.25 μM BFA was added for the final 12hrs. Antibody specific for KRT19 tail was used for detection. A one-way ANOVA with Turkey’s multiple comparison was performed. A representative experiment of three independent experiments is shown with two to three replicate wells per experiment. **(C-D)** Immunoblots are shown of total cellular lysates (**C**) or cell surface proteins **(B)** isolated from PDA FC1242 cells expressing doxycycline-inducible mSt or mSt-ATG4B^C74A^. **(E)** The fold-change in KRT8 and KRT19 secretion occurring with doxycycline induction of mSt-ATG4B^C74A^ is shown. A representative experiment of three independent experiments is shown. Two-way Anova with Sidaks multiple comparison test. Mean ± SEM is plotted; CTRL=control; ns= not significant, *p<0.05, ***p<0.001, ****p<0.0001.

Prompted by the observation that KRT8 is not biotinylated by Sec61-TurboID (**Figure 1I**) despite evidence that it interacted with KRT19 on the surfaces of PDA cells (**Figure 1A**; **Figure S1**), we examined the potential role of autophagy in KRT8 secretion by cell surface protein biotinylation with the cell-impermeable sulfo-NHS-LC-biotin followed by streptavidin-pulldown and immunoblot analysis. Doxycycline-induction of mSt-ATG4B^C74A^ was observed to inhibit autophagy by suppressing the conversion of LC3B-I to LC3B-II (**Figure 6C**), and cell surface biotinylation appropriately identified E-cadherin but not GAPDH. Consistent with the results of confocal microscopy and the ELISpot assays, biotinylation of KRT19 and KRT8 occurred (**Figure 6D**). However, the amounts of the biotinylated forms of these two cytoskeletal proteins were decreased by mSt-ATG4B^C74A^ induction (**Figure 6D-E**). We further explored the role of autophagy in keratin secretion using siRNA targeting *Beclin1* in Panc-1 cells. Here, siBeclin1 resulted in over a 90% reduction in Beclin1, but did not affect total KRT19 or KRT8 levels (**Figure S11D-E**). Similar to cells expressing mSt-ATG4B^C74A^, cell surface protein biotinylation of siBeclin1 transfected cells resulted in a decrease in surface KRT19 and KRT8 (**Figure S11F-G**). Thus, secretion of KRT19 and KRT8 is also regulated by secretory autophagy.

A recent report showed the protein translocon TMED10 can facilitate cytosolic protein entry into the ERGIC for autophagy dependent UcPS^27^. If Sec61-TurboID or ER-resident S1-10 transiently enters the ERGIC prior to being returned to the ER, and if KRT19 enters the ERGIC through TMED10, these assays may result in a false positive. To test this possibility, we used siRNA to knockdown TMED10 in Panc-1 Sec61-TurboID cells and assayed KRT19 biotinylation. siTMED10 transfection resulted in a robust reduction in TMED10, no change in total KRT19 within the cell, and a minor reduction in Sec61-TurboID expression (**Figure S12A-B**). However, this reduction in TMED10 did not affect KRT19 labeling by Sec61-TurboID. Furthermore, the Sec61-TurboID did not label ANXA1 (**Figure S3F**), a previously reported TMED10 substrate^27^. Together this suggest KRT19 enters the ER-Golgi network independent of TMED10.

### KRT19 Enters the ER in mouse PDA Tumors

Finally, to evaluate if KRT19 biotinylation occurs in PDA tumors, we seeded Panc-1 Sec61-TurboID tumors subcutaneously in nude mice. After tumors were palpable, mice were treated with or without 2 mM biotin containing drinking water for four days. Then the tumors were removed and homogenized, and the lysates were subjected to anti-KRT19 IP. In all analyzed tumors, immunoprecipitation of KRT19 and blotting with streptavidin-HRP revealed that when exogenous biotin was provided, KRT19 was biotinylated. Biotinylation CD44 served as the positive control (**Figure 7**). Therefore, KRT19 enters the ER in tumors formed *in vivo*.

**Figure 7:**
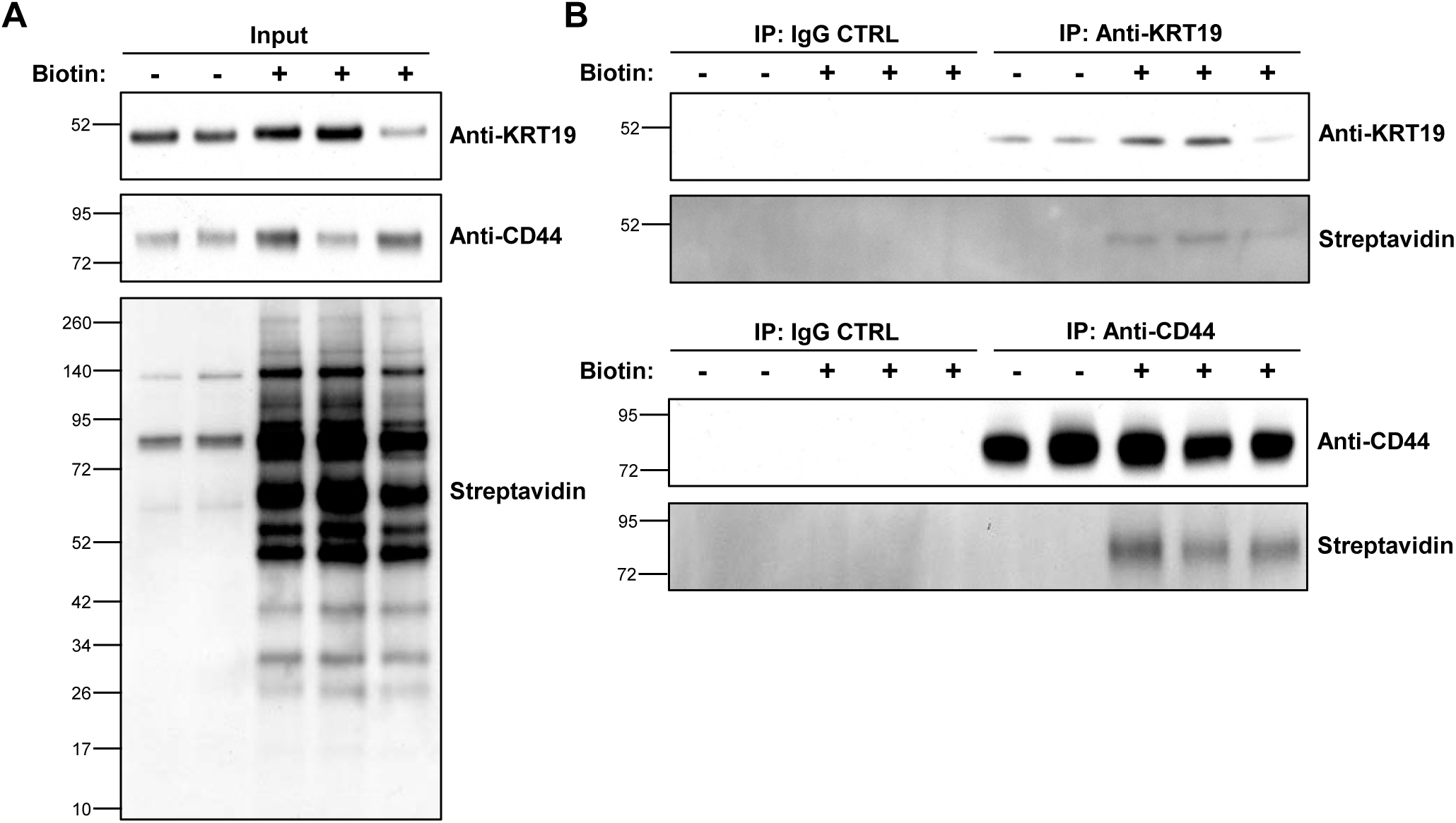
Entry of KRT19 into the ER in mouse PDA tumors. **(A-B)** Nude mice bearing subcutaneous tumors comprised of Panc-1 cells expressing Sec61-TurboID received plain drinking water (N=2) or drinking water containing 2 mM biotin (N=3) for four days. The tumors were removed and homogenized in RIPA buffer. The cleared lysates were incubated overnight with antibodies to KRT19 and CD44, respectively, or non-specific IgG. The cell lysates from the five tumors **(A)** and immunoprecipitated proteins **(B)** were applied to SDS-PAGE gels and the gels were immunoblotted with antibodies to KRT19 and CD44, and streptavidin-HRP, respectively.

## Discussion

The exclusion of T cells is proposed as a means by which carcinomas protect themselves from immune attack^42^. One mechanism that has been described in mouse PDA involves the inhibition of T cell motility by CXCL12 that is bound by KRT19 on cancer cells. The essential role for KRT19 in this process is shown by the T cell infiltration that occurs in tumors formed by PDA cells lacking expression of KRT19^3^. Furthermore, the incidence and number of KRT19-secreting cells has been correlated with overt metastasis and a poorer survival in patients with breast cancer^14^. However, the means by which KRT19 traffics to the cell surface to enable its T cell-exclusionary interaction with CXCL12 is unknown since KRT19 lacks a classical SP.

Here, we showed KRT19 is secreted through two pathways. First, KRT19 enters and is secreted from the ER. This occurs despite KRT19 lacking a canonical ER-targeting SP, apparently as the unique head domain of KRT19 functionally serves this purpose. Using two secretory system resident TurboID constructs and a split-GFP reporter, we have shown this is KRT19-specific function and does not occur with GAPDH, β-actin, β-tubulin, KRT8, or KRT18. Second, KRT19, and its binding partner, KRT8, can also be externalized via secretory autophagy. We propose that this circumstance enables the formation of KRT8/KRT19 microfilament-like network on the surface of PDA cells that was observed.

Canonical SPs lack a conserved consensus motif but are instead marked by a positively charged N-terminal region, a central hydrophobic region that transiently interacts with Sec61, and a C-terminal region containing the signal peptidase cleavage site^43^. When the SP of a nascent polypeptide exits the ribosome it engages with the SRP and this complex is delivered to the Sec61 translocon, priming it for SP insertion^44^. Here the hydrophobic region of the SP can interact with the transmembrane helices of Sec61 and access the lipid bilayer^16–19^. Despite the diversity of SP sequences, they engage the lateral gate helices of Sec61, suggesting they may do so via different molecular means during the translocation process. The head domain of KRT19 lacks this classic SP structure, but still interacts with both the SRP and Sec61. Whether KRT19 can engage the SRP and open Sec61 directly is unclear, and it may have assistance from an intermediate chaperone.

Why KRT19 lacks a canonical SP may be explained by its need to perform both intracellular and extracellular functions. Variations in SP strength can govern the nascent protein’s ability to be directed to and translocated into the ER, and this differential trafficking plays a role under both pathological^25^ and physiological conditions, with the inefficient translocation of the ER chaperone calreticulin being an example of the latter^45^. Here, replacement of the endogenous SP of calreticulin with a more efficient SP aberrantly affected its cytosolic role, but did not perturb its normal ER function^45^. Therefore, the presence of an atypical, and likely weak, SP sequence within KRT19 presumably allows it to exert both its cytosolic and extracellular functions.

An important finding of this study is the observation that KRT19, but not KRT8, enters the ER, but externalization of both proteins would be necessary because one predicts that KRT19, without KRT8, would be unstable because its inability to form the extracellular, intermediate filament-like KRT8/KRT19 heterodimeric complex^3^ (**Figure 1A**). Perhaps the cell must exclude KRT8 from the ER because its spontaneous polymerization with KRT19 would cause excessive ER stress and cytotoxicity. This hypothesis is supported by the observation that entry of aggregation prone α-synuclein into the ER in Parkinson disease models causes ER stress induction and cell death^25^.

Although the secretion of mature cytoplasmic KRT19 is hampered by its ability to heteropolymerize into intermediate filaments, keratin solubility is promoted via phosphorylation^46^, providing cytosolic keratins the ability to be externalized. Accordingly, Ju et al. (2015) showed that Akt phosphorylation of KRT19 at S35 helped solubilize KRT19 and facilitate its association with membranous HER2 in breast cancer cells^15^. S35 resides within the head domain of KRT19, and therefore this region may be important for both ER- and cytosolic-based secretion of KRT19. Furthermore, a recent publication by Zheng et al. (2022) highlighted the role of Sec61 in releasing calcium to specify ER-localized autophagosome initiation sites^47^. Therefore, while we studied how these pathways can individually influence KRT19 secretion, future studies should determine how autophagy may influence KRT19 entry into the ER or potentially ER to Golgi trafficking.

The role of autophagy was once thought to be that of bulk degradation, but the ability of autophagy to target specific proteins for degradation is becoming increasingly apparent^48^. Additionally, this degradative function of autophagy can be extended beyond roles in regulating protein homeostasis and cellular adaptation to nutrient stress to include the more nuanced functions in cancer where selective autophagy targets MHC-I for degradation and ultimate immune evasion^40^. Similar to degradative autophagy, our study adds to the growing body of literature that shows secretory autophagy is also a means by which cells modulate the immune response^49^ to mediate the secretion of KRT8 and KRT19, proteins that are required for immune exclusion of T cells in PDA.

Cancer often exploits normal physiology in its growth, and under physiological conditions the immune system must be regulated in areas of high foreign antigen flux, such as the gastrointestinal tract, to prevent aberrant immune activation. We suspect that at some sites subject to antigenic stress, such as the epithelial cells of small intestine and colon, CXCL12-coating is employed to suppress inappropriate immune damage. Indeed, the “coating” of colorectal cancer cells with CXCL12 has been shown in a clinical study to be involved in immune escape of this cancer^8^. While KRT19 is secreted via both conventional and unconventional means in PDA, other tumor or tissue types may overutilize the unconventional pathways. Determining if KRT19 is secreted under physiological conditions via the ER, secretory autophagy, or by both mechanisms would expand our understanding of both malignant and normal tissue immune regulation.

## Materials and Methods

### Cell Culture

The mouse PDA cancer FC1242 cell line was derived from LSL-KrasG12D/+; LSL-Trp53R172H/+; Pdx1-Cre (KPC) mouse tumors^28^. Panc-1 and HEK293T cells were procured from the CSHL cell line repository. All cells were cultured in DMEM (Corning, 10-013-CV) supplemented with 10% FBS (Seradigm, 1500-500), 100 U/mL penicillin, and 100 μg/mL streptomycin. Cell lines were periodically checked for mycoplasma contamination.

### Plasmid Construction

Sec61-TurboID and TM-TurboID were PCR amplified (NEB, M0492) from pCDNA5-Sec61b-V5-TurboID (Addgene, 166971) and pcDNA3-mem-TurboID (Addgene, 149409) respectively. These inserts were cloned into pLV-EF1a-IRES-Puro (Addgene, 85132) with the In-Fusion cloning system (Takara, 638948). *KRT18*, *KRT8*, and *KRT19*, and the associated truncations, were PCR amplified from Panc-1 cDNA. The S11 GFP fragment was added to the N- or C-terminus of the keratins by PCR, with a GSGS linker, and then cloned into pLV-EF1a-IRES-Puro with the In-Fusion cloning system. S1-10 was PCR amplified from FLAG-3C-sGFP1-10_pcDNA3.1(-) (Addgene, 182931), and a PDIA2 signal peptide and ER-retention tag was added by PCR to express S1-10 in the ER^33^. Removal of these exogenous sequences allowed for its expression in the cytosol. Cytosolic and ER-resident S1-10 was cloned into LV-EF1a-IRES-Blast, a gift from Dr. Christopher Vakoc, by In-Fusion cloning. pInducer-mSt and pInducer-mSt-ATG4B^C74A^ were gifts from Dr. Alec Kimmelman^40^. Plasmids were propagated in NEB Stable Competent *E. coli* (NEB, C3040H).

### Lentiviral Transduction

To generate Sec61-TurboID and TM-TurboID cell lines, Panc-1 and 1242 cells were transduced with lentivirus expressing either construct. The lentivirus was produced by cotransfecting a confluent 10 cm dish of HEK293T cells with 10 μg lentiviral vector together with the 7.5 μg of psPAX2 (Addgene, 12260) and 5 μg of pMD2.G (Addgene, 12259) with 80 μL 1 mg/mL of PEI (RnD, 7854). Virus was collected 36 and 48 hours post transfection. Cells were spin transduced at 1750 RPM for 45 minutes at room temperature with varying viral dilutions and 20 μg/mL polybrene (Sigma, TR-1003). The transduced cells were selected with 4 or 1 μg/mL puromycin (Sigma, P8833), respectively, for one week to generate stable cell lines. Cells transduced with a viral dilution to give a 30% infection rate were chosen. To generate FC1242 cells expressing mSt or mSt-ATG4B^C74A^, these constructs transduced with lentivirus using the aforementioned spin transduction. We selected cells over one week with 500 μg/mL G418 (Thermo, 10131035). Here, we used a 100% infection rate because the autophagy inhibition is through a dominant negative action ^40^. Cells were first sorted for mSt negative cells to remove cells with a leaky expression. Then the sorted cells were cultured with 1 μg/mL doxycycline (Takara, 631311) for 48 hours and then resorted for mSt positive cells. Individual clones were expanded.

### KRT19 IP for Silver Stain and Mass Spectrometry

Cells were grown in 10 cm dishes for 48 hours and then washed with PBS. 750 μL of RIPA buffer (1% Triton X-100, 0.1% SDS, 20 mM Tris pH 7.5, 150 mM NaCl) with protease and phosphatase inhibitor (Thermo, 78440) was added to the dish and the cells were scraped. Cell lysates were rotated at 4C for 30 minutes and then pelleted at 16,000xG for ten minutes at 4C. The protein concentration of the clarified lysate was measure with a BCA assay according to the manufacturers protocol (Thermo, A53225). Anti-KRT19 IP antibody (abcam, ab7755) or isotype control (Biolegend, 401407) were directly conjugated to epoxide dynabeads (Thermo, 14311D) according to the manufacturers protocol. 5 μg of conjugated antibody was then incubated with 500 μg of fresh FC1242 RIPA lysate overnight. The following day, the beads were washed three times with RIPA buffer, five minutes each. After the washes, the beads were transferred to a new tube. To prevent the disulfide bonds holding the antibody heavy and light chains from breaking, the IP was eluted with nonreducing 2X Laemmli buffer at 70C for five minutes. The entire IP product was loaded onto a 4-12% Bis-Tris SDS PAGE gel and run at 150V for 20 minutes and then 120V for one hour. The gel was then silver stained according to the manufacturers protocol (Bio-Rad, 161-0481).

### Panc1 and BXPC3 Sec61-BioID Secretome IP for Coomassie Stain and Mass Spectrometry

10 15-cm plates of parental or Sec61-BioID-expressing Panc1 and BXPC3 cells at 90% confluency were washed 3 times with PBS and incubated with 8 mL serum-free media containing 50 uM biotin for 24 hours. The media was collected and centrifuged at 500xg for 5min, then concentrated with 5K Protein Concentrators (Thermo, 88534). This concentrated media was then subjected to biotin and salt clean-up using Zeba Dye and Biotin Removal Spin Columns (Thermo, A44301). The media was then pre-cleared with 50 μL of Protein G Magnetic Beads (Thermo, 10003D) for 1 hr at room temperature while 50 μL Streptavidin C1 beads (Thermo, 65002) per sample were blocked for 1hr in 5% BSA in TBST at room temperature. The pre-cleared media was then incubated with the blocked streptavidin beads overnight at 4C. The beads were washed 6 times in TNET buffer (50 mM Tris-HCl pH 7.5, 150 mM NaCl, 5 mM EDTA, 1 % Triton X 100) prior to elution with 2X LDS (Thermo, NP0008) containing reducing agent (Thermo, NP0009) for 10min at 95C. The entire IP product was loaded onto a 4-12% Bis-Tris SDS PAGE gel and run at 130V for 30 minutes. The gel was then Coomassie stained according to the manufacturers protocol (Sigma, RSB).

### In-Gel Proteolysis Protocol

Silver stain solution was removed from each gel slice using 70 μL of 100mM sodium thiosulfate/30mM potassium hexacyanoferrate (III). Gel slices were washed 4x with 200 μL 50 mM ammonium bicarbonate (ABC)/33% acetonitrile (ACN) for five minutes at 55C under agitation. Gel bands were then dehydrated with 100 μL ACN. ACN was removed and gel bands were dehydrated by vacuum centrifugation for 10 minutes. Gel bands were reconstituted in 50 μL fresh 3 mM TCEP/50mM ABC and incubated at 55C for 10 minutes. The excess TCEP solution was removed. Gel bands dehydrated with 100 μL ACN until bands turned white. ACN was then removed. 50 μL fresh 10 mM CEMTS/ACN was added, and the tubes were incubated for 55C for 10 minutes. Excess CEMTS solution was then removed. Gel slices were washed once with 100 μL 50 mM ABC/33% ACN for 5 minutes at 55C under agitation. 100 μL of ACN was added and incubated for five minutes at 55C under agitation. ACN was removed and the gel bands were dehydrated by vacuum centrifugation. Next 100ng/μL of Trypsin (Sequencing grade modified porcine trypsin, Promega) was prepared in 100 mM ABC. 1000 ng (10 µl) of Trypsin to each band for five minutes on ice. 50 μL of 0.02% Protease Max surfactant was added to the gel bands in 50 mM ABC to cover the gel bands for 10 minutes on ice. The digest then continued at 37C overnight. The next day 50 μL of 80% ACN with 1% TFA was added to each gel bands and incubated for 15 minutes at 55C under agitation. Eluates were collected in clean tubes, which was repeated, pooling the eluates. Eluates were lyophilized by vacuum centrifugation. Peptide pellets were resuspended in 20uL of Loading Buffer (5% DMSO, 0.1% TFA in water). 2 μL of each sample was then injected.

### LC/MS QC and Analysis

For Co-IP experiments, peptides were loaded on a 20 cm x 75 μm ID column packed with Reprosil 1.9 μm C18 silica particles (Dr. Maischt). Peptides were resolved applying a segmented ACN gradient, consisting of a 4.5-30% ACN gradient in water (both mobile phases containing 0.1% formate) over 85 minutes, followed by 30-50% over 5 minutes, at constant flowrate of 200 nl/min. Eluting peptides were ionized by electrospray (constant 2200V) using silica emitters (4 cm length, 10 μm tip ID, FossilIonTech), mounted on a metal zero-dead volume union (50 μm, Valco) and transferred into the Lumos Orbitrap mass spectrometer (Thermo). The MS was operated in data-dependent mode and set to collect 120,000 resolution precursor scans (m/z 380-2000 Th) every 3 seconds. Targets for fragmentation were excluded if intensity was <1000 or if z<2 or or z>5, and dynamic exclusion was applied after one observation for 30 s (mass tolerance 10 ppm). Precursor ions selected for fragmentation were isolated using quadrupole (1.2 Th isolation window) and fragmented by HCD at stepped 30,35,40% normalized energy. Spectra were collected in the orbitrap at linear trap at unit resolution with maximum injection time set at 35 ms and AGC set to 200% (20,000 ions).

For secretome experiments, peptides were loaded via 10 cm x 100 μm ID trap column packed with 5 μm aqua C18 particles (Phenomenex) onto a 30 cm x 75 μm ID analytical column packed with Reprosil 1.9 μm C18 silica particles (Dr. Maischt), and resolved on an 80 minute 5-35% acetonitrile gradient in water (0.1% formic acid) at 200 nl/min. Eluting peptides were ionized by electrospray (2200V) and transferred into a Thermo Scientific Exploris 480 mass spectrometer. The MS was set to collect 120K resolution precursor scans (m/z 380-2000 Th). Precursor ions were selected from HCD fragmentation at stepped 28,33,38% normalized energy in data-dependent mode. Spectra were collected in the orbitrap at 15K resolution, with first mass locked to 100 Th.

Files were searched using the Mascot scoring function within Proteome Discoverer, with mass tolerance set at 5 ppm for MS1, and 0.3 Da for MS2. Spectra were matched against the UniProt mouse database and a database of common contaminants (cRAP). M-oxidation and N/Q-deamidation were set as variable modifications. Peptide-spectral matches were filtered to maintain FDR<1% using Percolator. Extracted ion chromatograms of precursor ions intensities were integrated for label-free quantification across samples, and used as quantitative metric for protein relative quantification. The mass spectrometry proteomics data have been deposited to the ProteomeXchange Consortium via the PRIDE^50^ partner repository with the dataset identifier PXD051583 and PXD057028.

For the secretome experiments, the Log_2_ fold-change cutoff of 3.5 for proteins considered as “hits” was determined by plotting the distribution of all fold-changes of the Sec61-BioID pulldown over the WT counterpart for the BXPC3 experiment. The distribution was bimodal, and Log_2_ fold-change of 3.5 was the local minimum between the two peaks in the distribution. This cutoff was applied to both the Panc1 and BXPC3 proteomics experiments.

### Tissue Culture Immunofluorescent Staining

Cells were grown in glass bottom plates (Chemglass, CLS-1812-024) until about 50-70% confluent, with or without a, 18 hour 50 μM biotin treatment. Cells were then washed with PBS and then fixed for ten minutes at room temperature with 4% PFA. Cells were washed once with PBS for five minutes, and then permeabilized for ten minutes at room temperature with 0.1% Triton X-100. Cells were washed once with PBS for five minutes, and then blocked with 1% BSA for one hour at room temperature. Primary antibody was incubated overnight in PBS. The following day, the cells were washed three times, five minutes each, with PBS. An appropriate fluorochrome conjugated secondary antibody was added at 2 μg/mL in PBS for one hour at room temperature. Simultaneously, the cells were stained with streptavidin-AF568 at 2 μg/mL in PBS (Thermo, S11226). The cells were then incubated with DAPI (Thermo, R37606) for ten minutes at room temperature and then washed twice with PBS for five minutes each. Cells were imaged using a Leica SP8 confocal microscope and the images were processed with the Leica LAS X suite (v3.7.4.23463).

### Cytosolic and ER Fractionation with Protein Extraction

The following protocol was adapted from Jagannathan et al. (2011)^32^. Panc-1 cells were grown until 70% confluent in a 10 cm dish and then washed with PBS. Cytosolic content was extracted with 1.5 mL of 0.015% digitonin solution for ten minutes on ice (0.015% digitonin (Sigma, CHR103), 1 mM MgCl_2_, 0.1 mM EGTA, 110 mM KCl, 25 mM HEPES pH 7.5, 1 mM fresh DTT, 40 units/mL murine RNAse inhibitor (NEB, M0314L), protease and phosphatase inhibitor (Thermo, 78440)). The cells were then quickly washed with 4 mL of 0.004% Digitonin buffer (same as above). ER content was then extracted with 1.5 mL of 2% DDM solution for ten minutes on ice (2% w/v DDM (Sigma, D4641-1G), 10 mM MgCl_2_, 200 mM KCl, 25 mM HEPES pH 7.5, 2 mM fresh DTT, 40 units/mL murine RNAse inhibitor (NEB, M0314L), protease and phosphatase inhibitor (Thermo, 78440)). If protein was to be analyzed, these products would be next clarified by centrifugation at 16,000xG for ten minutes at 4C and then quantified by BCA (Thermo, A53225).

### Streptavidin Pulldown of Sec61-TurboID Cell Lysate

Sec61-TurboID cells were plated in 10 cm dishes overnight and then treated with or without 50 μM biotin (Thermo, B20656) for 18 hours. Afterward, the cells were washed once with PBS and then lysed with 750 μL of RIPA buffer for 30 minutes at 4C and then pelleted at 16,000xG for ten minutes at 4C. The protein concentration of the clarified lysate was measure with a BCA assay according to the manufacturers protocol (Thermo, A53225). 250 μg of clarified lysate was incubated with 250 μL of a 50% NeutrAvidin agarose (Thermo, 29200) slurry in 300 μL of RIPA buffer with protease/phosphatase inhibitor for 30 minutes. The beads were then washed four times, two minutes each, with RIPA buffer with protease/phosphatase inhibitor, and the beads were eluted by boiling in Laemmli loading buffer.

### Immunoprecipitation of Sec61-TurboID and TM-TurboID Cell Lysate

Cells were plated in 10 cm dishes overnight and then treated with or without 50 μM biotin (Thermo, B20656) for the indicated times. Cell treatments include a vehicle control, 10 μM ESI (Tocris, 3922) for six or 17 hours, 50 μg/mL CHX (SelleckChem, S7418) for six or 24 hours, 100 nM or 250 nM CB-5083 (SelleckChem, S8101) for 18 hours. Afterward, the cells were washed once with PBS and then lysed with 750 μL of RIPA buffer for 30 minutes at 4C and then pelleted at 16,000xG for ten minutes at 4C. The protein concentration of the clarified lysate was measure with a BCA assay according to the manufacturers protocol (Thermo, A53225). 250 μg of clarified lysate was incubated with 2.5 μg of antibody in a total of 250 μL of RIPA buffer with protease/phosphatase inhibitor overnight at 4C. 20 μL of Protein G Dynabeads (Thermo, 10003D) per IP were blocked overnight at 4C with 1% BSA in RIPA buffer. The following day, the beads were washed once for five minutes with RIPA buffer and then incubated with the antigen-antibody complexes for two hours at 4C. The beads were them washed three times, five minutes each, with RIPA buffer with protease/phosphatase inhibitor. The IP was transferred to a new tube and the beads were eluted by boiling in Laemmli loading buffer.

### Split-GFP Transfection

HEK293T cells were grown in a 6-well plate until 95% confluent. After which, the medium of the cells was changed to fresh complete medium. 2.5 μg of both the S11 and S1-10 vectors were added to 250 μL of Opti-MEM (Thermo, 31985070), mixed and then combined with 40 μL of 1 mg/mL of PEI (RnD, 7854) for 15 minutes at room temperature. Then, this mixture was added dropwise to the cells. The medium was changed 24 hours after transfection, and then allowed to express for an additional 24 hours before the cells were analyzed by flow cytometry. Flow cytometry analysis was conducted using FlowJo (v10.8.1).

### Long Read Sequencing and Genome Mapping

RNA was isolated from FC1242 and Panc-1 cells (Qiagen, 74134) and cDNA was synthesized with Oligo(dT) primers. cDNA was sequenced on a PromethION P24 with a ONT SQK-PCS110 kit. The base called reads were aligned to the reference genome (hg38 for Panc1 and m39 for FC1242) using minimap2 (v2.26) with the parameter ‘-ax splice’. All primary alignments to KRT19 that have a mapping quality of at least 30 were extracted with ‘samtools view’. The coverage along the gene was visualized as a UCSC Genome Browser track. The long read sequencing can be accessed at SRA with the reference PRJNA1101971.

### Sec61β and SRP Immunoprecipitation

Cells were grown in 10 cm dishes for 48 hours and then washed with PBS. 750 μL of 1% Triton (1% Triton X-100, 20 mM Tris pH 7.5, 150 mM NaCl) with protease and phosphatase inhibitor (Thermo, 78440) was added to the dish and the cells were scraped. Cell lysates were rotated at 4C for 30 minutes and then pelleted at 16,000xG for ten minutes at 4C. The protein concentration of the clarified lysate was measure with a BCA assay according to the manufacturers protocol (Thermo, A53225). 500 μg of clarified lysate was incubated with 5 μg of antibody in a total of 500 μL of lysis buffer with protease/phosphatase inhibitor overnight at 4C. 35 μL of Protein G Dynabeads (Thermo, 10003D) per IP were blocked overnight at 4C with 1% BSA in lysis buffer. The following day, the beads were washed once for five minutes with lysis buffer and then incubated with the antigen-antibody complexes for two hours at 4C. The beads were them washed three times, five minutes each, with lysis buffer with protease/phosphatase inhibitor. The IP was transferred to a new tube and the beads were eluted by boiling in Laemmli loading buffer. For SRP IPs, 40 units/mL of Rnase inhibitor (NEB, M0314) was added to every step.

### Synthesis and Purification of Recombinant Human KRT19

Using Panc-1 cDNA as a template full-length and truncated KRT19 amplicons were produced by PCR using Q5 High-Fidelity master mix (NEB, K0492L). KRT19 constructs were either cloned into pGEX-4T1 (Addgene, 78827) with an N-terminal GST tag, or pET-28b(+) (EMD, 69865) with an N-terminal His tag. Vector and KRT19 inserts were inserted by restriction enzyme digests and subsequently ligated (NEB, M0202s). Stable competent E. coli (NEB, C3040) was transfected via heat shock with each plasmid, expanded, and sequenced to confirm appropriate construct incorporation. Sequenced plasmids were next transfected into BL21 cells (NEB, C2530H) and transfected BL21s were expanded in LB with ampicillin at 37C with shaking until the OD600 reached 0.4–0.8. Protein production was induced with 400 μM IPTG (Thermo, 15529019) at 16C overnight. Total proteins were isolated with B-PER (Thermo, 78248), and the insoluble pellets were resolubilized in 6 M urea in 50 mM Tris-HCl. The solubilized inclusion bodies in 6 M urea were buffer exchanged to 50 mM Tris HCl by Zeba Spin Desalting Columns (Thermo, 89883), then the supernatants were incubated with Glutathione Magnetic Agarose Beads (Thermo, 78602) to isolate GST tagged KRT19. For His-tagged KRT19s, His GraviTrap (Cytiva, 11003399) was used to isolate the proteins, then the eluted proteins were buffer exchanged in 50 mM Tris HCl buffer containing 200 mM guanidine. 10 ng of each protein was immunoblotted to determine the target epitope of each anti-KRT19 antibody.

### ELISpot Procedure and Quantification

The following protocol was adapted from Alix-Panabières et al. (2009)^14^. MSIP ELSIpot plate wells (Mabtech, 3654-TP-10) were activated with 20 μL of 35% ethanol for one minute and then washed five times with 200 μL of DI water. Then 100 μL capture antibody was added in PBS overnight at 4C. Wells were then washed three time with 200 μL of PBS and then blocked for two hours with 200 μL of 20% FBS/PBS at 37C. After, the wells were washed once with 200 μL of PBS and then 1000 Panc-1 cells or 1000 or 1500 FC1242 cells were plated in 200 μL of complete culture medium. Before treatment, cells were allowed to adhere to the wells for six hours, after which, 20 μL of the vehicle or treatment chemical was added to the wells in a 10X concentration for a final 1X concentration. Panc-1 cells were treated with a final concentration of 10 μM BFA (Thermo, 00-4506-51) for 17 hours, 10 μM GCA (SelleckChem, S7266) for 12 hours, 10 μM ESI (Tocris, 3922) for 17 hours, and 50 μg/mL CHX (SelleckChem, S7418) for 24 hours. FC1242 cells expressing Dox-inducible mSt or mSt-ATG4B^C74A^ were treated with 1 μg/mL doxycycline (Takara, 631311) for a total of 48 hours, with or without the final 12 hours also consisting of a 1.25 μM BFA treatment. Afterwards, the cells were washed away with three washes of 200 μL of 0.2% PBST and then five 200 μL PBS washes. Next, 100 μL of detection antibody in 1% BSA/PBS was incubated overnight at 4C. The following day the wells were washed five times with 200 μL of PBS. 100 μL of 1/1000 anti-rabbit-HRP (Biolegend, 406401) in 1% BSA/PBS was incubated per well for one hour at room temperature. Next, the wells were washed five times with 200 μL of PBS. 100 μL of filtered ELISpot TMB (Mabtech, 3651-10) was added per well for five to seven minutes, and then decanted. The wells were washed five times with 200 μL of DI water and let dry overnight in the dark.

As no ELSIpot plate reader was available, we 3D printed punches that could punch of strips of ELSIpot membranes from the MSIP plates. We first applied clear tape to the back of the wells, so that the membranes would be captured when punched out. These membranes were then imaged with a Nikon SMZ1500 microscope. Images were imported into Fiji (v1.54g), converted to black and white 16-bit images, inverted, and the region of interest was segmented. To quantify the ELISpot puncta number and size, the ComDet plugin (v0.5.5) was used.

### CellTiter-Glo Assay

Cells were plated in 96-well assay plates (Agilent, 204626-100) and treated identically as with the ELISPOT assays. Afterwards, 150 μL of medium was removed leaving the cells and 50 μL of medium. Then 50 μL of CTG reagent (Promega, G9241) was added per well and incubated according to the manufacture’s protocol.

### Cell Surface Protein Biotinylation and Isolation

FC1242 cells were cultured to 90% confluency in 6 cm plates. The cells were washed once with PBS, and then 1 mL of 0.5 mg/mL Sulfo-NHS-LC-Biotin (Thermo, A39257) in PBS was added to the dish for 10 minutes at room temperature. After, the plates were washed twice with 5 mL of cold Tris-buffered saline (Thermo, 28376), and then the cells were harvested by scraping in 10 mL of the cold Tris-buffered saline with protease/phosphatase inhibitor mixture (Thermo, 78445). The cells were pelleted and lysed with 250 μL of cold RIPA buffer with protease/phosphatase inhibitor. The protein concentration was quantified by a BCA (Thermo, A53225); 250 μg of the lysate was incubated with 250 μL of a 50% NeutrAvidin agarose (Thermo, 29200) slurry in 300 μL of RIPA buffer with protease/phosphatase inhibitor for 30 minutes. The beads were them washed four times, two minutes each, with RIPA buffer with protease/phosphatase inhibitor, and the beads were eluted by boiling in Laemmli buffer.

### siRNA Mediated Knockdown

Panc1 Sec61-TurboID or WT FC1242 cells were plated in 6-well plates and were then transfected at ∼40% confluency with 50 nM of the indicated siRNA (Dharmacon, D-001810-10-20; L-055895-00-0010; L-003718-00-0010) in 2 mL of serum-free medium with 5 μL of DharmaFECT 1 (Dharmacon, T-2001-03) for 18 hours. The cell medium was then changed to 2 mL of complete media to recover for an additional 24 hours prior to cell surface labeling (FC1242) or 18 hours of biotin labeling (Sec61-TurboID Panc1).

### Animal Tumor Engraftment, Treatment, and Harvesting

Nude mice seven weeks of age were purchased from Jackson Laboratory (Jackson, 002019). After receiving, the mice were housed in a Helicobacter-free room for 2 weeks before each experiment. Panc-1 cells expressing Sec61-TurboID were trypsinized and resuspended in sterile PBS at 10,000,000 cells/mL. Before injection, the right flank of the acclimated Nude mice was shaved and then 1,000,000 cells were injected into the right flank under anesthesia with isoflurane. After the tumor was palpable, the mice were given 2 mM biotin containing drinking water for four days (Thermo, B20656). The mice were then sacrificed via carbon dioxide inhalation and then cervical dislocation. The excised tumors were then snap frozen in liquid nitrogen. 100 mg of each tumor was homogenized (Qiagen, 9002756) in 1 mL of RIPA buffer with protease/phosphatase inhibitor mixture (Thermo, 78445) and incubated at 4C for 30 minutes and then pelleted at 16,000xG for ten minutes at 4C. The protein concentration of the clarified lysate was measure with a BCA assay according to the manufacturers protocol (Thermo, A53225). 500 μg of clarified lysate was incubated with 5 μg of antibody in a total of 500 μL of RIPA buffer with protease/phosphatase inhibitor overnight at 4C. 35 μL of Protein G Dynabeads (Thermo, 10003D) per IP were blocked overnight at 4C with 1% BSA in RIPA buffer. The following day, the beads were washed once for five minutes with RIPA buffer and then incubated with the antigen-antibody complexes for two hours at 4C. The beads were them washed three times, five minutes each, with RIPA buffer with protease/phosphatase inhibitor. The IP was transferred to a new tube and the beads were eluted by boiling in Laemmli loading buffer. All animal experiments were approved by the Cold Spring Harbor Laboratory (CSHL) Institutional Animal Care and Use Committee in accordance with the NIH Guide for the Care and Use of Laboratory Animals.

### Quantification and Statistical Analysis Software

Immunoblots were quantified using Fiji (v1.54g). Graphs were generated in, and statistical analysis was conducted using, GraphPad Prism 10 (v10.0.2).

### Materials Availability

All unique/stable reagents generated in this study are available from the Lead Contact with a completed Materials Transfer Agreement.

## Acknowledgments

We thank Dr. Alec Kimmelman and Dr. Christopher Vakoc for providing lentiviral vectors. We also acknowledge the contributions of the CSHL Animal, Microscopy, Flow Cytometry, NGS, and Mass Spectrometry cores and the CSHL Machine Shop. We thank Dhivyaa Anandan for assisting with manuscript editing. This work was supported by a Lustgarten Foundation Distinguished Scholar Award (D.T.F.), the Thompson Family Foundation (D.T.F.), the Cedar Hill Foundation (D.T.F.), the National Cancer Institute (D.T.F.), NIH T32 Training Grant T32GM008444 (P.M.), the NIH NCI F30 Ruth L. Kirschstein National Research Service Award (P.M.), The Lustgarten Foundation (J.P.K; D.A.T.) and the NIH Cancer Center Support Grant 5P30CA045508 (J.P.K; D.A.T.). Schematics were created with Biorender.com.

