## Supplemental Figures for "Signal peptide-independent secretion of keratin-19 by pancreatic cancer cells"

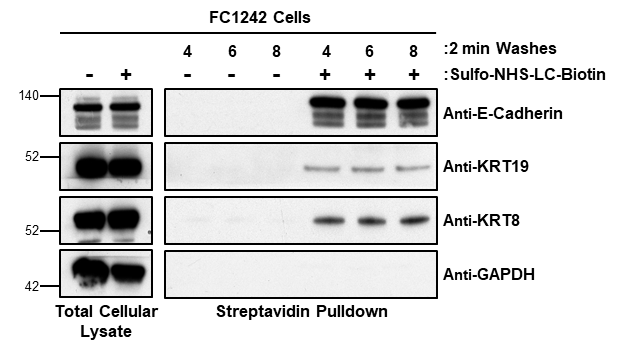
**Figure S1:** The externalization of KRT8 and KRT19 by mouse PDA cells. FC1242 PDA cells were surface-labeled with sulfo-NHS-LC-biotin, or not. Lysates were prepared in RIPA buffer and biotin-labeled proteins were isolated with bead-immobilized streptavidin. The beads were subjected to variable number of two-minute washes in RIPA buffer before elution in 2.5x Laemmle buffer. The total lysates and biotinylated proteins were analyzed by SDS-PAGE and immunoblotting with antibodies to E-cadherin, KRT19, KRT8, and GAPDH. Shown are representative results of three replicate experiments.

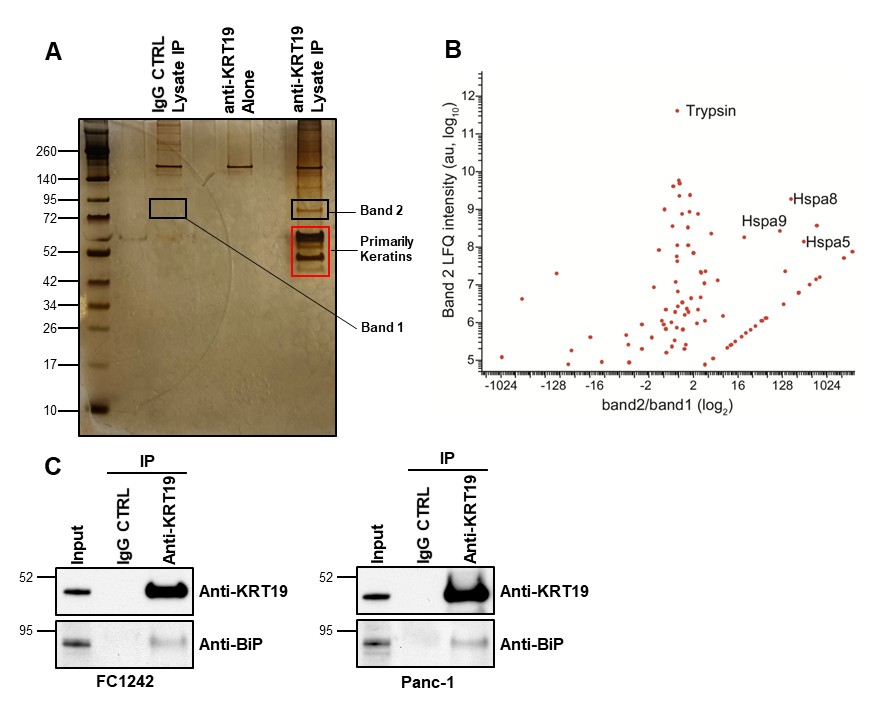

**Figure S2:** Co-immunoprecipitation of KRT19 and BiP. **(A)** Silver stain is shown of immunoprecipitates obtained with IgG CTRL and anti-KRT19 antibody from RIPA lysates of mouse FC1242 PDA cells. Bands 1 and 2 were excised and analyzed by LC-MS. **(B)** Volcano plot of peptide spectra isolated in Bands 1 and 2. Experiment conducted once. **(C)** The RIPA lysates of FC1242 and Panc-1 cells were immunoprecipitated by IgG CTRL or anti-KRT19 antibody and subjected to SDS-PAGE and immunoblotting with anti-KRT19 and anti-BiP antibodies. Shown are representative results of three replicate experiments.

\

**Table S1:** LC-MS identified anti-KRT19 antibody co-immunoprecipitating proteins.

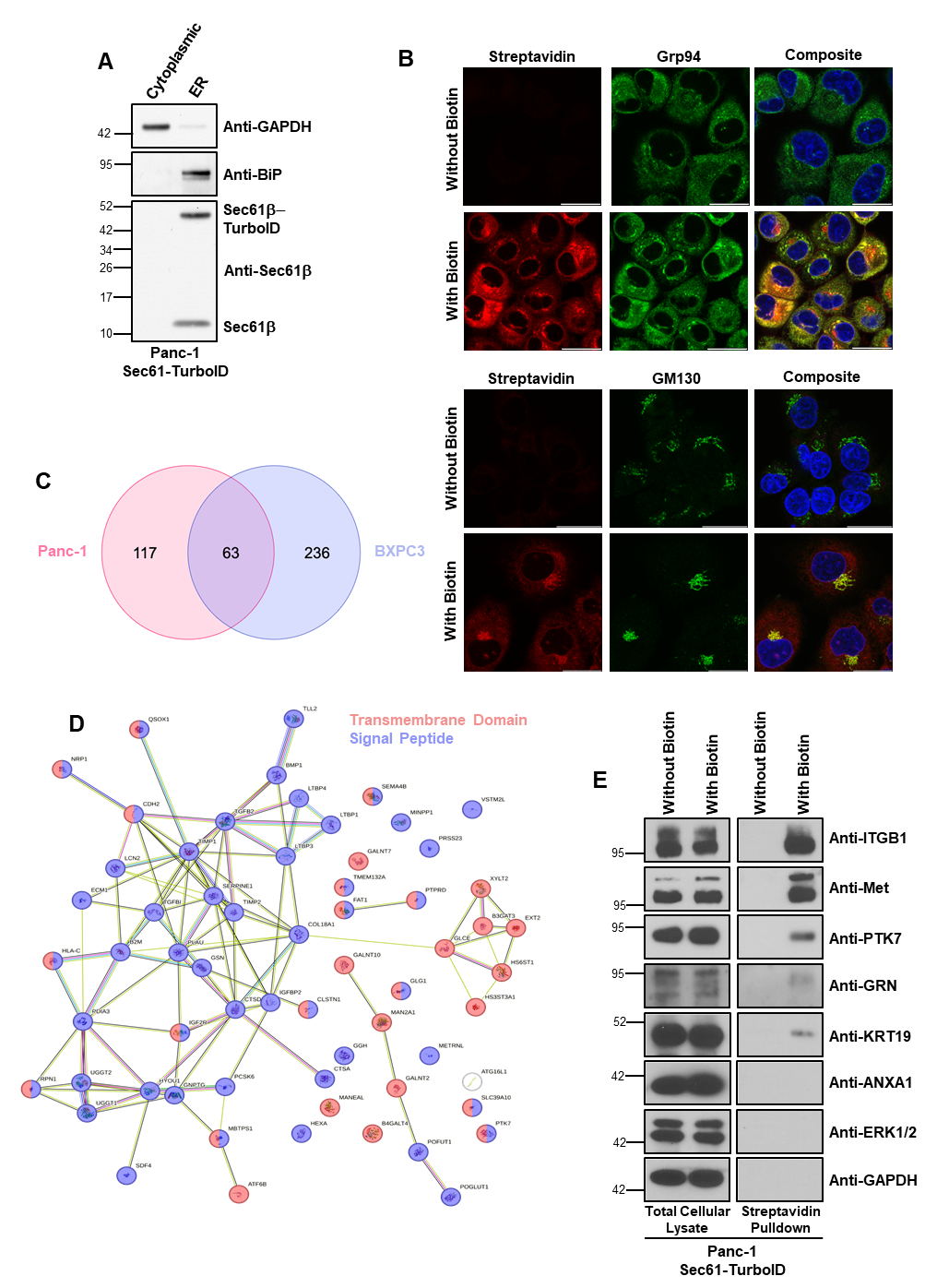
**Figure S3:** Sec61-TurboID restriction to the ER compartment. **(A)** Panc-1 cells expressing Sec61-TurboID were sequentially fractionated with digitonin to isolate cytoplasmic proteins, and DDM to isolate the ER resident proteins. Cytoplasmic and ER fractions were analyzed by immunoblot. Shown are representative results of three replicate experiments. **(B)** Panc-1 cells expressing Sec61-TurboID were cultured for 18 hrs in the presence or absence of 50 μM biotin. After fixation and permeabilization, the cells were stained with fluorochrome-conjugated streptavidin and anti-Grp94 antibody or streptavidin and anti-GM130 antibody to show biotin incorporation into the secretory pathway. Shown are representative results of three replicate experiments. (Scale bar = 10 μm). **(C)** BXPC3 and Panc-1 cells expressing Sec61-TurboID and their parental counterparts were cultured for 18 hrs in the presence of 50 μM biotin in serum-free DMEM. The media from these cells was incubated with streptavidin beads to isolate biotinylated proteins. The proteins were eluted, subjected to SDS-PAGE, silver staining, and analyzed by LC-MS. The Venn diagram depicts the number of proteins identified as enriched by LC-MS (Log_2_ fold change > 3.5) in each Sec61-TurboID expressing cell line compared to the parental cell line. **(D)** STRING-DB plot depicting the overlapping proteins from the LC-MS experiment, with proteins containing a signal peptide colored in blue and those with a transmembrane domain colored in red. **(E)** Panc-1 Sec61-TurboID cells were cultured in the presence or absence of 50 μM biotin for 18hrs and then lysed with RIPA buffer. The cell lysates were incubated with streptavidin beads to isolate biotinylated proteins. Total cellular lysates and streptavidin-pulldowns were analyzed by immunoblots.

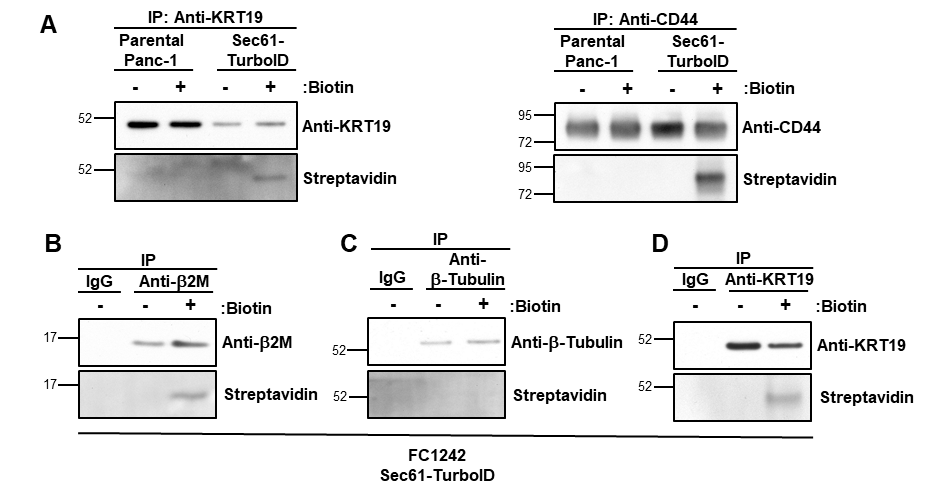

**Figure S4:** KRT19 entry into the ER in mouse FC1242 PDA cells. **(A)** Parental or Sec61-TurboID-expressing Panc-1 cells were cultured for 18 hrs in the presence or absence of 50 μM biotin and lysed with RIPA buffer. The lysates were incubated with antibodies to KRT19 and CD44, respectively, or non-specific IgG control. Immunoprecipitates were obtained and analyzed by immunoblot for biotin incorporation. Shown are representative results of three replicate experiments. **(B-D)** FC1242 cells expressing Sec61-TurboID were cultured for 18 hrs in the presence or absence of 50 μM biotin. The cells were lysed with RIPA buffer, and lysates were incubated overnight with antibody to β2M **(B)**, β-Tubulin **(C)**, and KRT19 **(D)**, respectively, or IgG control. The immunoprecipitated proteins were analyzed by immunoblot for biotin incorporation. Shown are representative results of three replicate experiments.

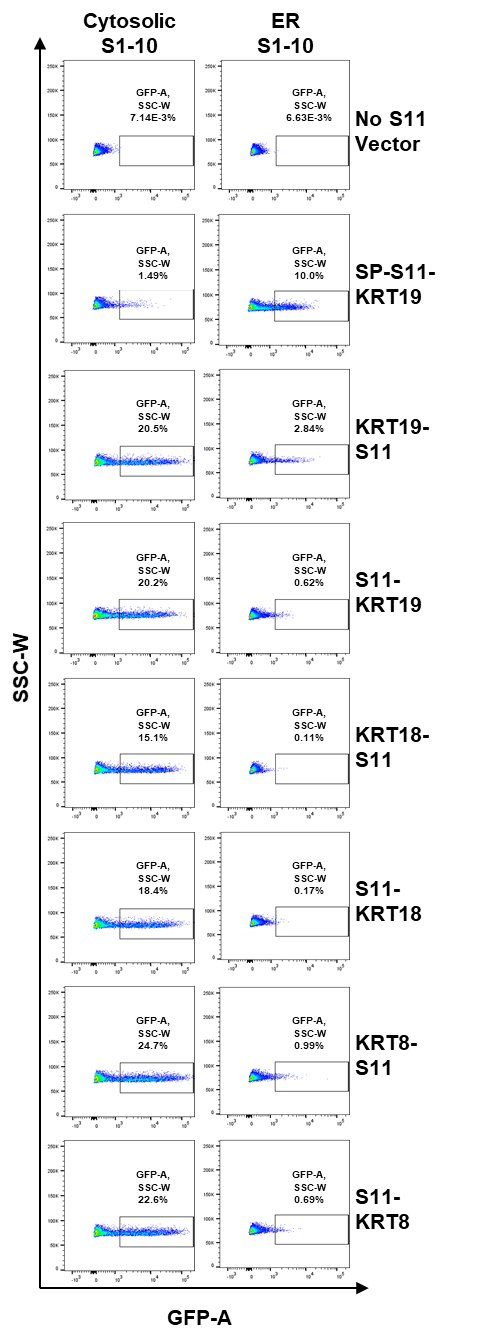

**Figure S5:** The Split-GFP assay and identification of ER-resident KRT19. Flow cytometry plots of HEK293T cells transfected with ER-directed or cytosolic GFP-domains S1-10, and KRT8, KRT18 and KRT19, respectively, with their N- or C-terminal fused to the GFP-S11 domain. A SP-bearing KRT19 was used as a positive control for ER entry. Shown are representative results of three replicate experiments.

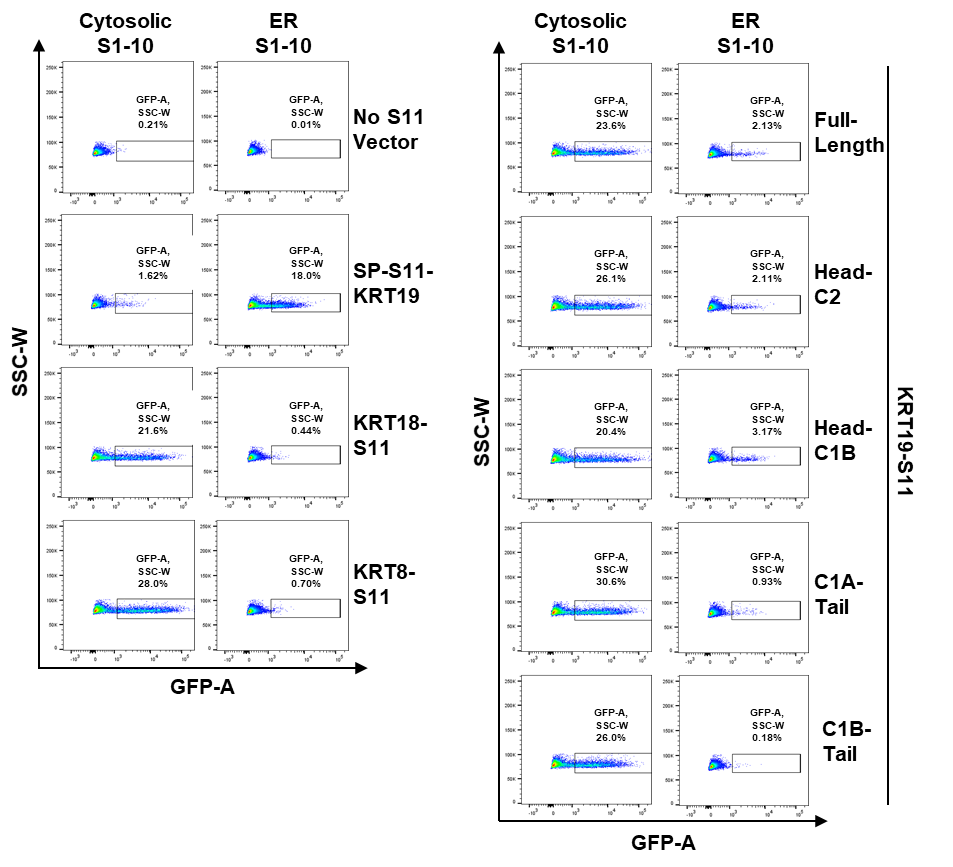

**Figure S6:** The head domain of KRT19 and entry into the ER. Flow cytometry plots of HEK293T cells transfected with ER-directed or cytosolic GFP-domains S1-10, and KRT19 fragments fused to the C-terminal GFP-S11 domain. A SP-bearing KRT19 was used as a positive control for ER entry. Shown are representative results of three replicate experiments.

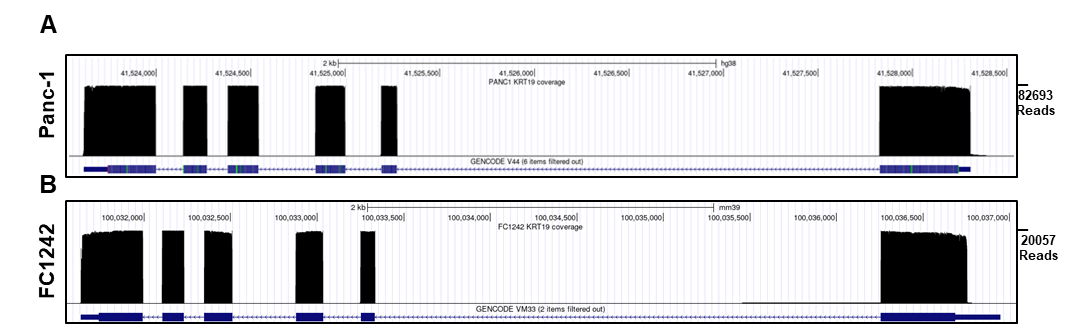

**Figure S7:** Germline encoding of *KRT19* transcripts. **(A-B)** Long-read sequencing of Panc-1 cells **(A)** and FC1242 cells **(B)** was performed. The *KRT19* transcripts were mapped to the respective *KRT19* genomic loci. Bottom blue tracts indicate *KRT19* intronic and exonic regions. Upper black track indicates the coverage of mapped transcripts along the *KRT19* gene. Experiment conducted once.

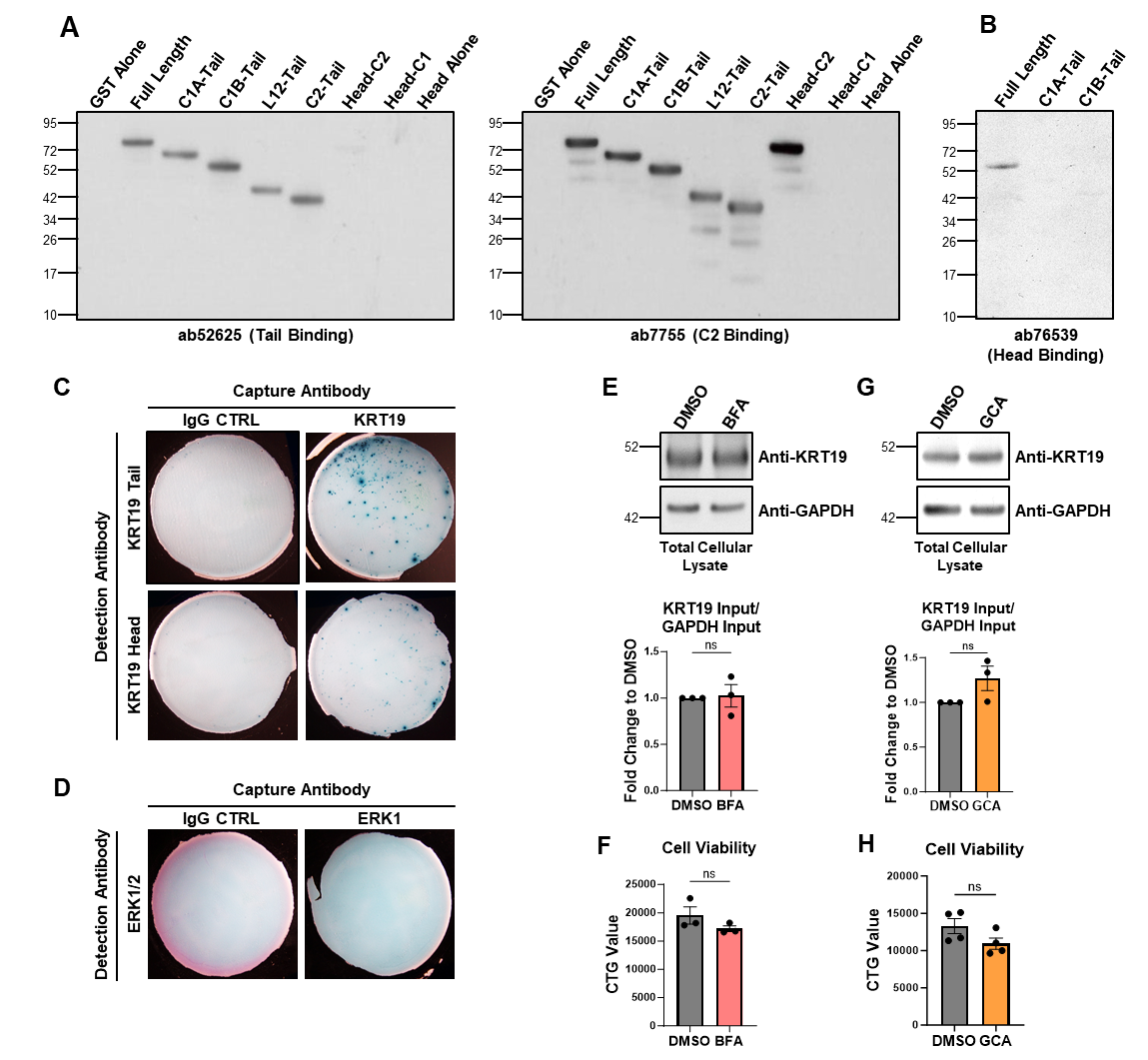

**Figure S8:** Secretion of KRT19 from Panc-1 cells. **(A)** The binding sites of ab52625 and ab7755 were validated by immunoblotting GST-tagged recombinant KRT19 truncations. **(B)** The binding site of ab76539 was validated by immunoblotting His-tagged recombinant KRT19 truncations. Shown are representative results of three replicate experiments. **(C-D)** Panc-1 cells, 1000, were cultured for 48 hrs in ELISpot wells coated the following capture antibodies: IgG CTRL, anti-KRT19 **(C)**, or anti-ERK1 **(D)**. One of two anti-KRT19 **(C)** or anti-ERK1/2 **(D)** antibodies were used for detection. Shown are representative results of two replicate experiments. **(E-F)** Panc-1 cells were cultured for 16 hrs with DMSO or 11 μM BFA. Total cell lysate GAPDH and KRT19, respectively, were analyzed by immunoblot (one representative of three independent experiments shown) **(E).** Cell viability was evaluated by a CTG assay (one representative of three independent experiments is shown) **(F)**. **(G-H)** Panc-1 cells were cultured for 12 hrs in the presence of DMSO or 10 μM GCA. Total cellular GAPDH and KRT19 was measured by immunoblots (one representative of three independent experiments is shown) **(G)** or cell viability was evaluated by a CTG assay (one representative of three independent experiments is shown) **(H)**. Welch’s T test **(E/G)** or a Student’s T test **(F/H)** was performed. Mean ± SEM is plotted; ns=not significant; CTRL=control

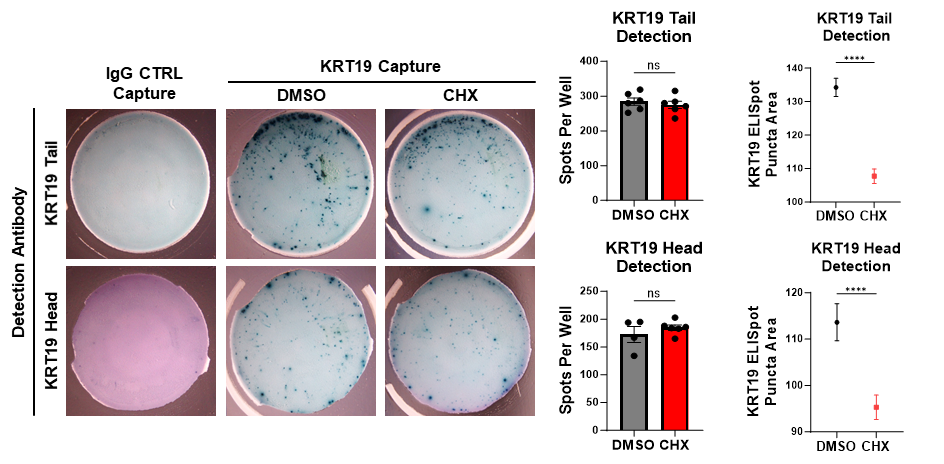

**Figure S9:** Co-translational secretion of KRT19. Panc-1 cells, 1000, were plated in ELISPOT wells coated with IgG CTRL or anti-KRT19 as capture antibodies. The cells were cultured for 24 hrs in the presence of DMSO or 50 μg/mL CHX. Antibodies to the KRT19 head or tail domains were used for detection. A Students T test was performed on the spots per well and puncta area. Shown are representative results of two replicate experiments with two to three replicate wells per experiment. Mean ± SEM is plotted; ns=not significant; CTRL=control; ****p<0.0001

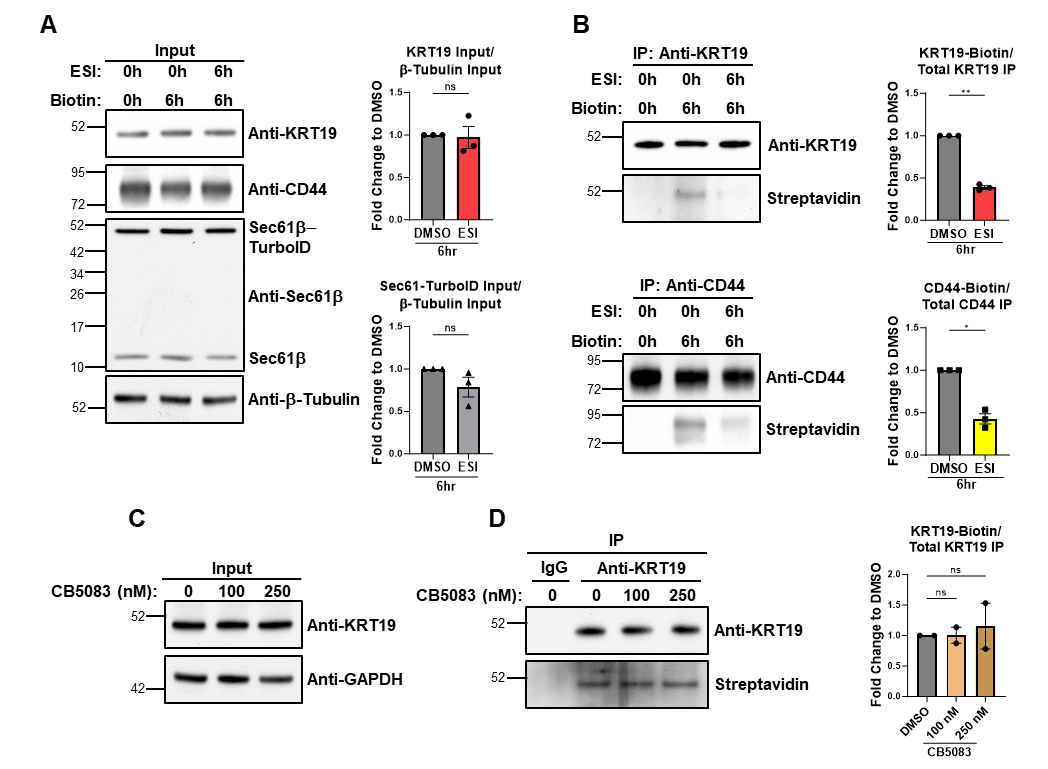

**Figure S10:** Entry of KRT19 into the *ER* *via* Sec61. **(A-B)** Panc-1 expressing Sec61-TurboID cells were cultured for 6hrs with 10 μM ESI, in the presence or absence of 50 μM biotin. The cells lysed in RIPA buffer, and lysates were incubated overnight with antibodies to KRT19 and CD44, respectively, or IgG control. Total lysates **(A)** and the immunoprecipitated proteins **(B)** were analyzed by immunoblots. One presentative of three independent experiments is shown. Welch’s T test was performed. **(C-D)** Panc-1 cells expressing Sec61-TurboID were cultured for 18 hrs with DMSO or CB5083, in the presence of 50 μM biotin. The cells lysed with RIPA buffer and lysates were incubated overnight with antibodies to KRT19 and CD44, respectively, or non-specific IgG control. Total lysates **(C)** and immunoprecipitated proteins **(D)** were analyzed by immunoblots. One representative of two independent experiments is shown. A one-way ANOVA (Brown-Forsythe test) with Dunnett's multiple comparisons was performed. Mean ± SEM is plotted; h=hours; ns=not significant, *p<0.05, **p<0.01.

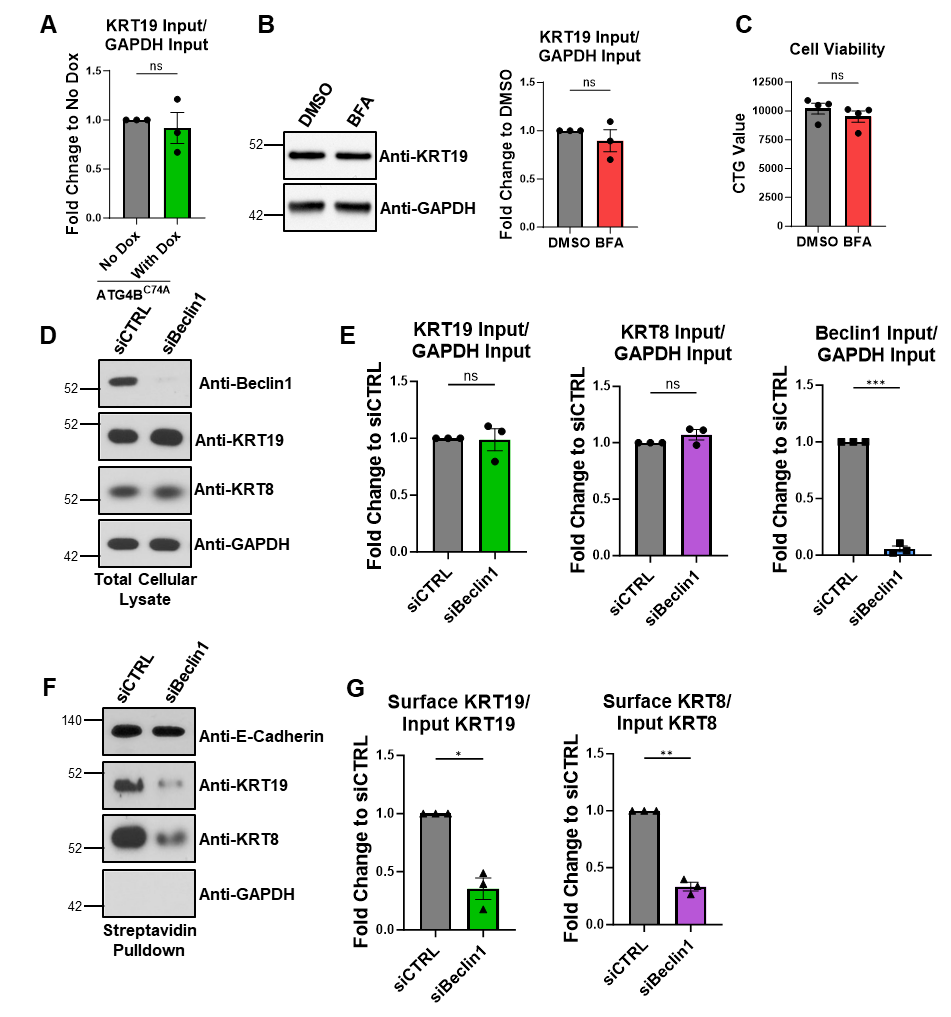

**Figure S11:** Secretion of KRT19 and KRT8 via autophagy. **(A)** Mouse PDA FC1242 cells expressing doxycycline-inducible mSt-ATG4B^C74A^ were cultured for 48 hrs in the presence or absence of 1 μg/mL Dox. Cell lysates were prepared with RIPA buffer and KRT19 and GAPDH in the cell lysates was analyzed by immunoblots. One representative of three independent experiments is shown. **(B-C)** Mouse PDA FC1242 cells expressing doxycycline-inducible mSt-ATG4B^C74A^ were cultured for 12 hrs in the presence of DMSO or 1.25 μM BFA. Cell lysates were prepared with RIPA buffer and KRT19 and GAPDH in the cell lysates was analyzed by immunoblots. One representative of three independent experiments is shown. **(B)** Cell viability was assayed by a CTG assay. One representative of three independent experiments is shown **(C)**. Welch’s T test **(A/B)** or a Student’s T test **(C)** was performed. **(D-G)** Immunoblots are shown of total cellular lysates (**D**) or cell surface proteins **(F)** isolated from PDA FC1242 cells transfected with siCTRL or siBeclin1. **(E)** The fold-change in total cellular KRT19, KRT8, and Beclin1 with Beclin1 knockdown. **(G)** The fold-change in cell surface KRT19 and KRT8 with Beclin1 knockdown. A representative experiment of three independent experiments is shown. Welch’s T test was performed. Mean ± SEM is plotted; CTRL=control; ns= not significant, *p<0.05, **p<0.01.

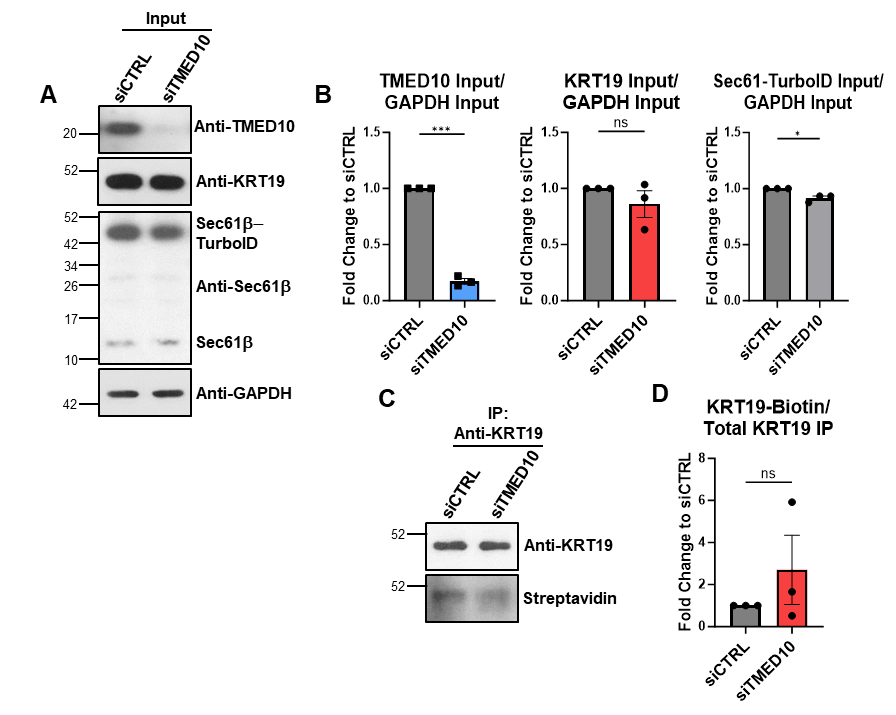

**Figure S12:** Influence of TMED10 on KRT19 ER-entry. Panc-1 expressing Sec61-TurboID cells were transfected with siCTRL or siTMED10, in the presence or absence of 50 μM biotin. The cells lysed in RIPA buffer, and lysates were incubated overnight with antibodies to KRT19 or IgG control. Total lysates **(A)** and the immunoprecipitated proteins **(C)** were analyzed by immunoblots. **(B)** The fold-change in total cellular TMED10, KRT19, and Sec61-TurboID with TMED10 knockdown. **(G)** The fold-change in biotin-KRT19 relative to total KRT19 IP with TMED10 knockdown. A representative experiment of three independent experiments is shown. Welch’s T test was performed. Mean ± SEM is plotted; CTRL=control; ns= not significant, *p<0.05, ***p<0.001.

**Table S1: LC-MS identified anti-KRT19 antibody co-immunoprecipitating proteins.**

**Table S2:** Antibodies used in this study.

| **ANTIBODY** | **SOURCE** | **IDENTIFIER** |
| --- | --- | --- |
| Mm anti-KRT19 | abcam | ab7755 |
| Mm IgG1 Isotype Control | Biolegend | 401407 |
| Rb anti-KRT19 | abcam | ab52625 |
| Mm anti-BiP | Proteintech | 66574-1-Ig |
| Mm anti-GAPDH | Cell Signaling | 97166 |
| Rb anti-Sec61β | Proteintech | 15087-1-AP |
| Rb IgG Isotype Control | Cell Signaling | 3900 |
| Rb anti-Grp94-CL488 | Proteintech | CL488-14700 |
| Mm anti-GM130 | BD | 610822 |
| Mm anti-CD44 | Invitrogen | MA5-13890 |
| Rb anti- β2M | Cell Signaling | 12851 |
| Rb anti-KRT8 | abcam | ab53280 |
| Mm anti- β-Actin | Cell Signaling | 4967S |
| Rb anti-Biotin | Cell Signaling | 5597 |
| Mm anti- β-Tubulin | Thermo | MA5-16308 |
| Rb anti- β2M | Proteintech | 13511-1-AP |
| Rb anti-KRT19 | abcam | ab76539 |
| Mm anti-ERK1 | Thermo | 13-8600 |
| Rb anti-ERK1/2 | Cell Signaling | 4695 |
| Rb anti-SRP68 | Proteintech | 11585-1-AP |
| Rb anti-SRP19 | Proteintech | 16033-1-AP |
| Rb anti-V5 | Cell Signaling | 13202S |
| Rb anti-ITGB1 | Cell Signaling | 4706S |
| Rb anti-Met | Cell Signaling | 8198S |
| Rb anti-PTK7 | Cell Signaling | 25618S |
| Mm anti-GRN | Thermo | MA1-187 |
| Rb anti-Beclin1 | Cell Signaling | 3738S |
| Mm anti-TMED10 | Proteintech | 67876-1-Ig |
| Rb anti-ANXA1 | Cell Signaling | 32934S |
| Rb anti-RFP | Rockland | 600-401-379 |
| Rb anti-LC3B | Cell Signaling | 2775 |
| Mm anti-E-Cadherin | Cell Signaling | 14472 |
| Rb anti-KRT19-AF568 | abcam | ab203445 |
| Rb anti-KRT8-AF647 | abcam | ab192468 |
| Dk anti-rabbit-IgG-HRP | Biolegend | 406401 |
| Goat anti-mouse-IgG-HRP | Biolegend | 405306 |
| Veriblot | abcam | ab131366 |
